# Rationalised experiment design for parameter estimation with sensitivity clustering

**DOI:** 10.1101/2023.10.11.561860

**Authors:** Harsh Chhajer, Rahul Roy

## Abstract

Quantitative experiments are essential for investigating, uncovering and confirming our understanding of complex systems, necessitating the use of effective and robust experimental designs. Despite generally outperforming other approaches, the broader adoption of model-based design of experiments (MBDoE) has been hindered by oversimplified assumptions and computational overhead. To address this, we present PARameter SEnsitivity Clustering (PARSEC), an MBDoE framework that identifies informative measurable combinations through parameter sensitivity (PS) clustering. We combined PARSEC with a new variant of Approximate Bayesian Computation for rapid, automated assessment and ranking of designs. By inherent design, PARSEC can take into account experimental restrictions and parameter variability. We show that PARSEC improves parameter estimation for two different types of biological models. Importantly, PARSEC can determine the optimal sample size for information gain, which we show correlates well with the optimal number of PS clusters. This supports our rationale for PARSEC and demonstrates the potential to harness both model structure and system behaviour to efficiently navigate the experiment design space.

## Introduction

The scientific approach relies on careful experiments to understand complex systems. Experiments must not only provide quantitative information such as model parameter estimates but also be economically and practically viable. Accurate parameter estimation is essential for refining model representations and enhancing model predictions. Compared to traditional model-free techniques, Model-Based Design of Experiments (MBDoE) utilises system-specific models to optimise experiment design for improved parameter estimation [1, 2, 3, 4, 5, 6, 7]. MBDoE allows for a more effective evaluation of the experimental design space and offers a more comprehensive understanding of the variables and the interactions between the components. It can facilitate resource optimisation, decrease the number of trials, and improve the accuracy of the estimation of the model parameters [3, 4, 5] and model selection [6, 7]. Over the years, MBDoE has become widely applied in a variety of fields, including engineering and biomedicine, due to the increasing use of quantitative models to understand complex systems. Despite the advantages of MBDoE, its application in parameter estimation is hindered by the increasing intricacies of modern experiments and their inherent constraints.

Fisher’s Information Matrix (FIM) is a popular framework for designing experiments aimed at estimating model parameter values [3, 4, 5, 6, 7, 8]. It is based on the concept of the expected information gain from an experiment, which is calculated by taking the expected value of the second derivative of the log-likelihood function with respect to the model parameters [9, 10]. If changes in the values of the parameters have a major effect on the output of the model, it implies that the output is useful in providing information about the parameters. Nevertheless, FIM-MBDoEs usually assume a linear statistical model that links measurements to parameter values, requiring the linearisation of nonlinear models at expected ground-truth parameter values. Therefore, the efficiency of designs is markedly diminished when the guesses are not precise or accurate [9]. This limits the ability of FIM methods to incorporate parameter uncertainty in the experimental design. The complexity of matrix inversion and the need for a large experimental sample size to achieve the desired accuracy and a Gaussian noise distribution present additional difficulties that restrict the use of FIM-MBDoE [11]. Recently, new approaches have been proposed to address some of these issues [8, 12, 11, 10, 13, 14, 9]. For example, design space discretization can be employed to handle non-linear models, and the average information content and risk can be calculated for a large uncertainty to identify robust and reliable designs [14]. Additionally, the pseudo-inverse [15, 16, 17] can be used to tackle ill-conditioned matrices [8].

Alternately, Bayesian approaches to MBDoE operate independently of assumptions related to linearity or direct likelihood estimations [18, 19, 20, 21, 22, 23, 24, 25]. These techniques can take into account any prior information or convictions about the parameters and offer more flexibility that successfully sidesteps the issues with a single-point estimation. However, they suffer from high computational demands, particularly when dealing with complex systems or large design spaces, and often produce biased and potentially suboptimal designs when tailored to prespecified sample sizes [18, 22, 21, 20]. To address these limitations, certain algorithms employ a sequential greedy approach, which build on existing designs to manage complexity and improve design efficiency [26, 8, 19]. However, such design search approaches are influenced by the initial design choice and may be susceptible to local optima artefacts.

Here, we introduce the PARSEC DoE algorithm, which uses the model architecture of the system through parameter sensitivity analysis to direct the search for informative experiment designs. PARSEC recognises combinations of measurables with a low overlap in the vectors that represent their parameter sensitivity indices (PSI). These PSI indices indicate how the value of the measurable changes when the value of the parameter changes. We demonstrate that the precision of parameter estimates increases when the overlap in the PSI vectors is reduced, thus enabling PARSEC to generate an ‘optimal’ DoE effectively. We show that fuzzy c-means clustering of PSI can reduce the number of evaluations required to identify ‘optimal’ designs substantially. PARSEC combines the PSI evaluated at different parameter values to identify generalist and robust designs informed by the parameter sensitivities while accommodating the uncertainty in prior knowledge. PARSEC combines concepts from FIM-based and Bayesian MBDoEs to determine the best experimental designs. We find that the optimal number of PSI clusters is in agreement with designs that yield the highest information gain, thus introducing a new way to calculate the sampling frequency.

In order to implement PARSEC, a reliable high-throughput parameter estimation framework is needed to assess the predicted designs. Existing methods for parameter estimation are restricted by their inherent assumptions or are not suitable for high-throughput analysis. For instance, likelihood-based techniques are limited to systems where the likelihood function can be specified and a Gaussian distribution for parameter distribution and measurement noise can be assumed.[27, 28, 29, 30]). Rather than using error thresholds or correlated sampling, Approximate Bayesian Computation (ABC) based methods [30, 31, 32, 33]) typically rely on data-dependent stipulations. This can make the estimation vulnerable to local minima and initial biases. To alleviate this, here we develop the Approximate Bayesian Computation - Fixed Acceptance Rate (ABC-FAR) method [34] for parameter estimation for PARSEC designs. Our ABC-based algorithm uses a global and relative parameter sampling rejection criterion (FAR). Thus, ABC-FAR avoids data-dependent stipulations, making it suitable for automated analysis and fair comparison of estimation derived from different data sets. We show that parameter estimation via ABC-FAR is accurate, free of computational artefacts, and less susceptible to noise in the data and initial guess bias.

## Results

### Sensitivity driven design of experiments

The PARSEC (PARameter SEnsitivity driven Clustering based DoE) framework represents a MBDoE rooted in four key ideas (Figure 1). Firstly, it employs the parameter sensitivity of a variable as an indicator of its informativeness towards the estimation of a parameter value [35, 10]. The information content profile of a measurement is, thus, approximated using a vector of its parameter sensitivity indices (PSI) (Step 1, Figure 1). Secondly, PARSEC computes the PSI vectors at various parameter values that sample the distribution linked to parameter uncertainty. Concatenating the PSI vectors for a measurement candidate yields the composite PARSEC-PSI vector. This approach accounts for the uncertainty and helps inform robust DoE. The intricacies of creating the PARSEC-PSI vector are detailed in the Supplementary text (SI S2, Supplementary Figure S1). The third idea revolves around the selection of measurements in a manner that minimizes the overlap amongst the corresponding PARSEC-PSI vectors. This maximizes the overall information yield from their combination. The process involves clustering measurements based on their PARSEC-PSI vectors and subsequently selecting a representative measurement from each cluster (Step 2, Figure 1). Finally, we expect the performance of a PSI-clustering-based design to depend on the quality of clustering, which in turn hinges on the number of clusters. As the number of clusters dictates the size of the experimental sample, we posit that optimal clustering can provide a well-informed guess for the sample size, which can be tuned to identify efficient designs that are both economical and informative (Step 3, Figure 1). The choice of clustering depends on the design specifications. For predetermined sample sizes, k-means and c-means clustering algorithms are suitable, especially for PARSEC-PSI vectors which would typically be high-dimensional, continuous data. In this study, we focus on two alternate implementations of PARSEC, namely PARSEC(k) and PARSEC(c), that uses the k-means and c-means clustering algorithms respectively (see Methods).

**Figure 1:**
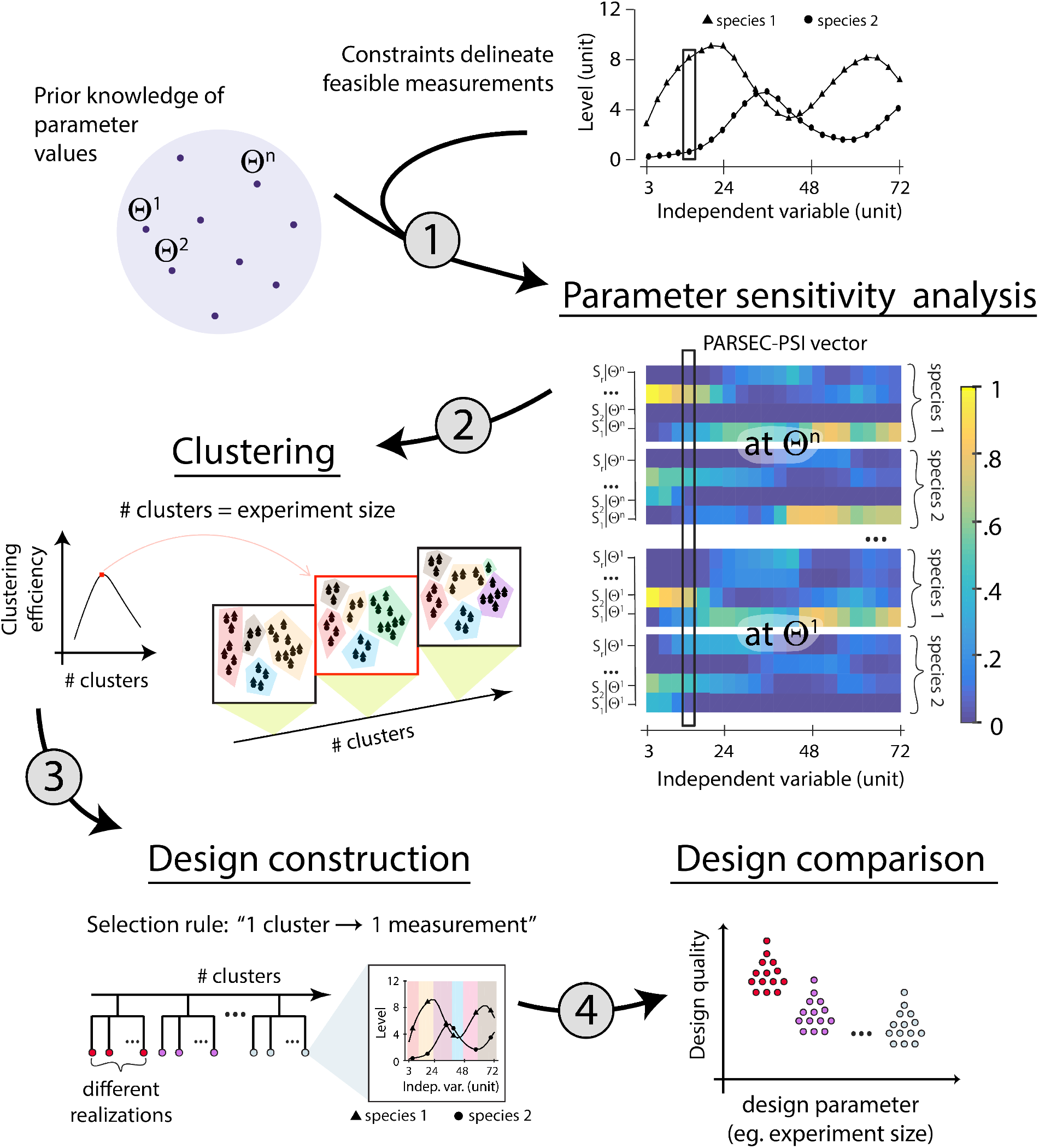
Schematic for PARSEC: PARameter-SEnsitivity driven Clustering based experiment design algorithm. Given the parameter uncertainty, PARSEC samples some representative values (Θ^*k*^) and evaluates parameter sensitivity indices (PSI) for each of the feasible measurement candidates there (Step 1). Here we assume that simultaneous measurement of species 1 and 2 constitute a single measurement candidate. The PSI vectors for the same measurement, but evaluated at different parameter values, are vectorially conjoined to create the PARSEC-PSI vector (enclosed in the black rectangle). Based on the similarity among the PARSEC-PSI vectors, the measurement candidates are clustered (step 2). One candidate from each cluster is randomly selected to constitute the predicted design (step 3). Thus the number of clusters used in partitioning translates to the experiment sample size, which can be optimized within the practical constraints of the experiment. Since clustering and subsequent candidate selection are stochastic, PARSEC delivers multiple realizations of design. The predicted designs are computationally compared to identify the optimal one (step 4). It should be noted that PSI vectors can come from continuous range and hence an appropriate clustering algorithm should be used. Furthermore, the set of candidates before clustering can be filtered optionally to enrich for high overall PSI values.

Optimizing the design parameters requires evaluation of the associated designs predicted. Moreover, given the stochastic nature of PSI-clustering and subsequent measurement selection, PARSEC predicts multiple designs. Thus the process of design comparison is vital for selecting the most favorable design from the array of designs predicted by the PARSEC framework (Step 4, Figure 1). In this context, we introduce the Approximate Bayesian Computation - Fixed Acceptance Rate (ABC-FAR) technique, designed to automate parameter estimation and facilitate impartial comparison of the generated experimental designs.

For a likelihood-free approach and wider applicability, we propose an ABC-based algorithm for parameter estimation. Like the other ABC methods, ABC-FAR iteratively refines the parameter value distribution using χ^2^ statistics. In each iteration (Figure 2a), it samples the current distribution estimate (step 2, Figure 2a), and among them chooses some based on the corresponding χ^2^ values to update the distribution estimate (steps 3 & 4, Figure 2a). However, ABC-FAR differs from other ABC methods in that it selects a fixed fraction (FAR) of parameter values with the lowest χ^2^ values to update the marginals. The rejection criterion employs relative comparison and eliminates the need for specifying absolute χ^2^ thresholds, thereby avoiding data-specific customization. Unlike other relative rejection criterion-based approaches that involve local comparisons with neighboring parameter combinations, ABC-FAR performs a global comparison, considering all combinations sampled from the marginal together. Latin Hypercube Sampling (LHS, [36, 37]) is employed to sample the marginal, eliminating the need for transition kernels. Additionally, a small noise is introduced in the sampling distribution, resembling Simulated Annealing [38], to enhance robustness against local minima and initial guess biases (step 1, Figure 2a).

**Figure 2:**
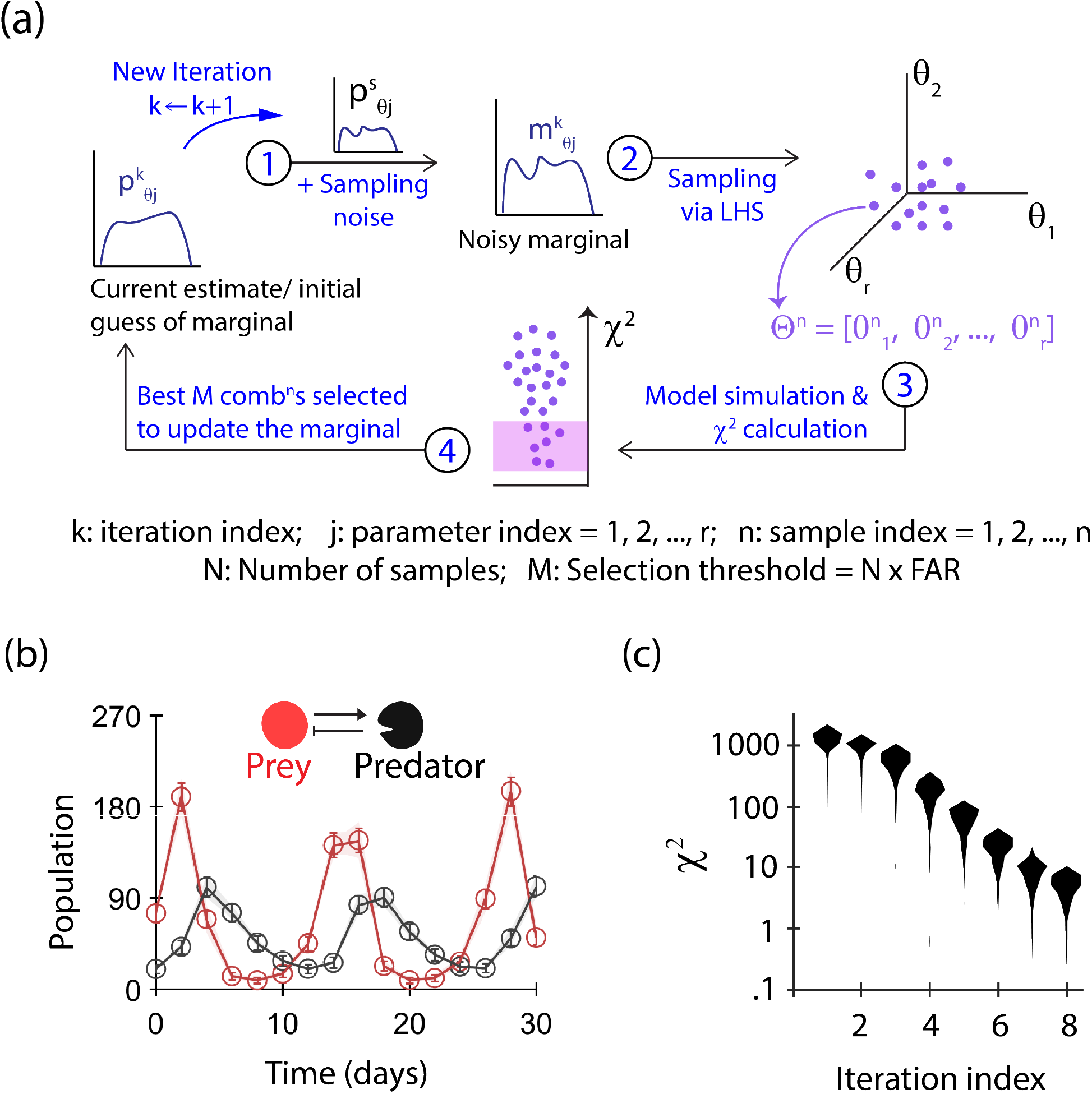
Schematic and performance of ABC-FAR. (a) Schematic: ABC-FAR uses an iterative, ABC approach to estimate the likely distribution of model parameter values that minimizes the χ^2^ function (see Methods). In an iteration, ABC-FAR (1) augments the current guess of marginal with a sampling noise, (2) uses them to sample N parameter combinations (via Latin Hypercube Sampling [36]), (3) at which the χ^2^ function is evaluated, (4) to select M combinations (=FAR × N) with the lowest χ^2^ values to update the marginal, and the iteration restarts with the updated estimate of the marginal. (b) Model fitting using ABC-FAR: We use ABC-FAR to fit the Lotka Volterra model to a computationally generated data set. The figure shows the model prediction due to each of the 250 parameter combinations (thin solid lines) selected after eight iterations, along with the averaged dynamics (thick solid lines) and the data set used for fitting (open circles). Here we used a FAR value of 0.25 and the history-dependent update strategy (see Methods). (c) ABC-FAR reduces χ^2^ iteratively: The distribution of χ^2^ values corresponding to the parameter combinations selected in each of the eight iterations are shown.

### Accurate and robust parameter estimation using ABC-FAR

We demonstrate the performance of ABC-FAR using the popular Lotka-Volterra model. We simulate the model for a known combination of parameter values (ground truth) and initial conditions to generate synthetic data (Figure 2b). Assuming the same initial conditions, we use ABC-FAR to estimate parameter values using this data. Estimation is done in the logarithmic scale (base 10) of parameter values to explore a large dynamical range. We use a uniform initial guess for all parameters to indicate a lack of prior knowledge about the values. We see that the deviation between model predictions and data (χ^2^ statistics) decreases with every iteration of ABC-FAR, and the model predictions recapitulate the data well (Figure 2b, c). The algorithm iteratively transforms the prior guess for the model parameters into sharp distribution around the corresponding ground truth value used (Figure 2c, Supplementary Figure S2). However, it doesn’t alter the marginal of a dummy parameter (Figure 3c), a parameter that doesn’t affect the model predictions (and hence χ^2^) but undergoes the same treatment as the model parameters. This suggests that the algorithm doesn’t introduce unwanted computation artifacts.

**Figure 3:**
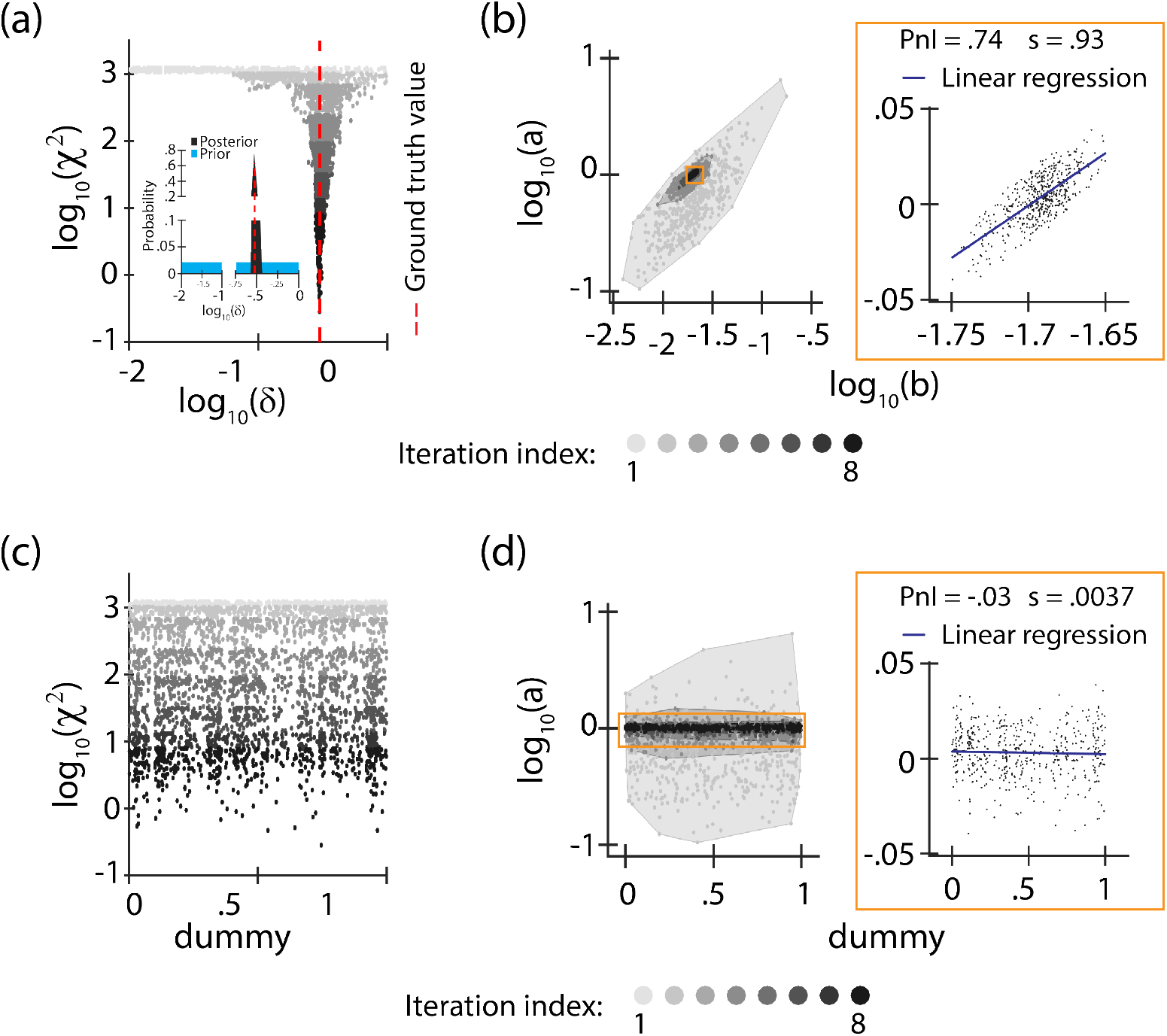
Accuracy and efficiency of the parameter estimation algorithm. (a) We plot the value of one of the model parameters (δ, death rate of predators) along with the χ^2^ value corresponding to the parameter combinations selected in every iteration associated with figure 2b. The inset show the initial guess and the estimated posterior for δ. The distribution converges to sharp distribution around the ground truth value. (b) The value of two model parameters (a: birth rate of prey, and b: prey death rate due to predation) corresponding to the selected combinations are shown. The correlation is used to approximate the practical non-identifiability, and the regression (inset) is used to measure the degree of compensation for the two model parameters. (c) We plot the value of a dummy parameters along with the χ^2^ value corresponding to the parameter combinations selected in every iteration associated with figure 2b. The absence of any structure in the selected dummy parameter values suggests an absence of computational artifacts due to the algorithm. (d) The value of a and the dummy parameter corresponding to the selected combinations are shown. The correlation and regression (inset) measures associated with dummy parameter, can be used to identify thresholds for statistical significance.

Additionally, we identify higher-order relations among model parameter values, conditioned on data. For example, the selected values of the ‘birth rate of prey’ (a) and ‘death rate of prey due to predation’ (b) show a strong positive correlation (PnI = .74, Figure 3b), indicative of significant practical nonidentifiability (PnI). The slope of linear regression (=0.93) suggests that changes in χ^2^ due to a 1% increase in log_10_(a) value can be compensated by .93% increase in the value log_10_(b). Although local, the findings are statistically significant as realized by comparing them to similar analysis on dummy parameter (Figure 3d, Supplementary Figures S3 and S4). We further verify that the estimation is robust to measurement noise (Supplementary Figure S6) and see that ABC-FAR can easily be adapted to exploit the model structure and system-specific knowledge to speed up and improve convergence (Supplementary text SI S7, Supplementary Figure S7).

The efficiency of ABC-based methods typically depends on the implementation of their rejection criterion. In the previous analysis, we used a FAR value of 0.25. Here we vary it to see the effect on the algorithm’s efficiency. For this, we monitor two performance statistics at every iteration for the different implementations - the accuracy of estimation (inversely related to χ^2^ statistics) and cumulative computational cost (∝ EAR^*−*1^, described in methods, see Supplementary Figures S8, S9 and S10). The χ^2^ statistics monotonically decrease with iteration irrespective of the FAR value used; this is expected as we use the ‘history-dependent update strategy’ (strategy described in Methods). However, the estimation converges at higher accuracy (lower χ^2^ statistics) as the FAR value decreases. Among those reaching a particular level of accuracy, the schemes with higher FAR values (weaker rejection criteria) seem to be more efficient. The efficiency of our schemes is comparable to that of the popular ABC-SMC algorithm [32] (Supplementary Figure S10, comparison detailed in Methods).

### Identifying informative designs using PARSEC with ABC-FAR

We use PARSEC(k) (PARSEC employing k-means clustering, see Methods) and ABC-FAR to design experiments characterizing the parameters of a three-gene repressilator model (Elowitz and Leibler in 2000, details in Methods and SI). First, we establish the methodology with predetermined initial conditions and sample size, assuming an accurate prior knowledge of parameters. We account for practical considerations, such as (a) limiting the experiment duration, (b) preferring simultaneous measurements when multiple variables are observed, and (c) constraining the measurables based on feasibility (see Methods).

To validate PARSEC(k)’s performance, we compare PARSEC(k) designs (PD_*BC*_) against random designs (RD_*BC*_), for parameter estimation accuracy. The suffix indicates the variables being measured. We also evaluate the importance of clustering, using anti-PARSEC(k) designs (WD_*BC*_) where all measurements are selected from the fewest of clusters (the ones with the most candidates). The parameter estimation error for the random designs (RD_*BC*_ and RD_*AB*_) are approximately uniformly distributed (Figure 4a). Interestingly, the spread of errors becomes strongly skewed when we consider designs based on PSI vector clustering (PD_*BC*_ and WD_*BC*_, Figure 4a). The error for designs where all the measurements belong to a single cluster (WD_*BC*_) tends to be higher; whereas designs (PD_*BC*_) generated using measurements from different clusters are highly informative with lower error (barring few outliers). The top ten percentile of designs monitoring the levels of proteins B and C (RD_*BC*_, WD_*BC*_ and PD_*BC*_) are predominantly PARSEC-predicted ones (PD_*BC*_). Despite no significant bias in the selection of measurements in PD_*BC*_ compared to that of RD_*BC*_ (Figure 4c), the average estimation error for the PD_*BC*_ is about three times lower than that for the RD_*BC*_ (Figure 4a). Furthermore, the clustering-based algorithm samples from a small and informative sub-space of the experiment design space, indicated by the low variance in estimation error (Figure 4a) and performance of sub-samplings of PD_*BC*_ compared to that of RD_*BC*_ (Figure 4b).

**Figure 4:**
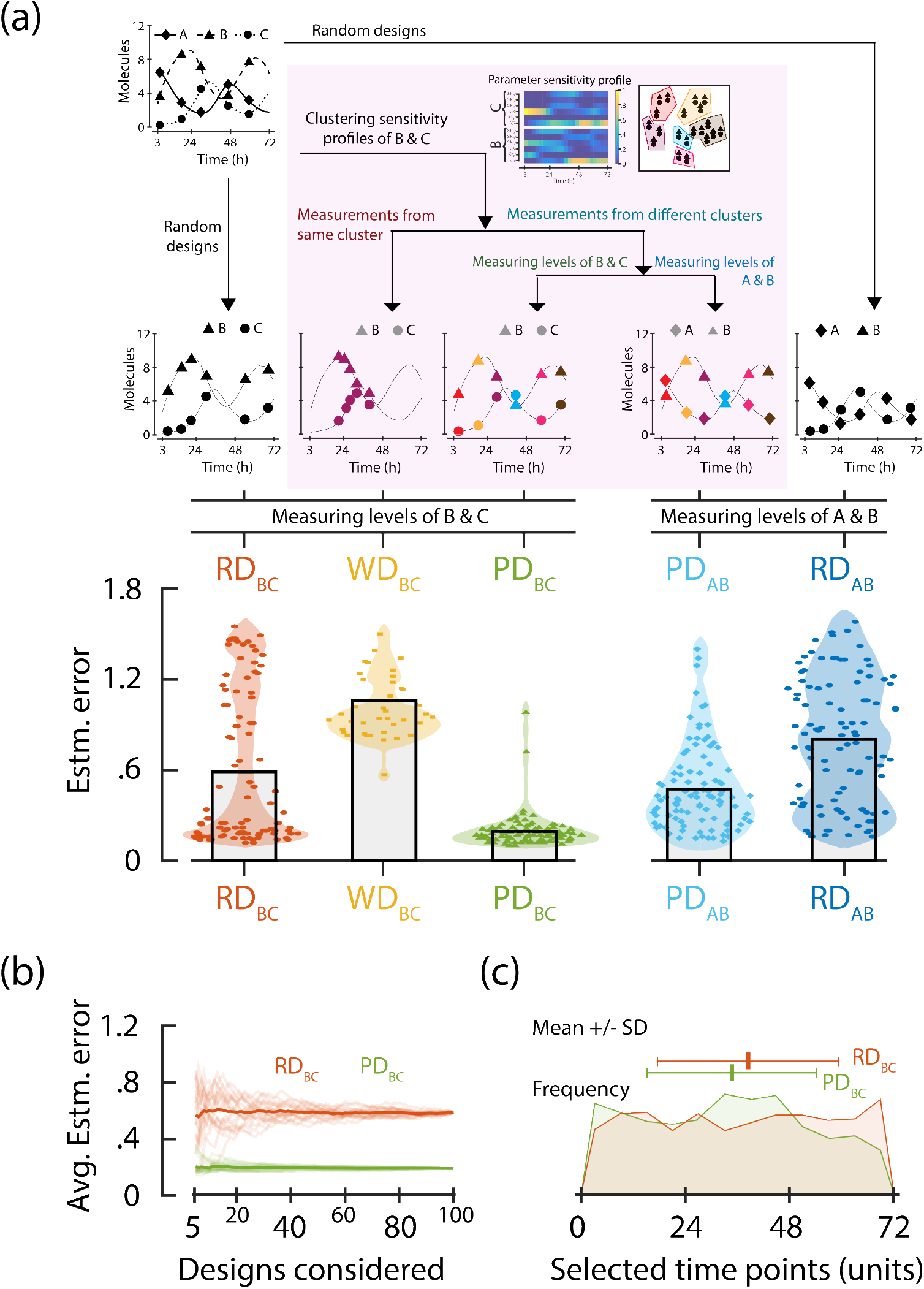
Performance of PARSEC-predicted designs. (a) To validate the performance of PARSEC(k), we compare designs generated by multiple modalities (illustrated in the top panel). RD_*BC*_ and RD_*AB*_ denote random designs where the measurement time points were chosen in an unbiased, non-repetitive manner. PD_*BC*_, WD_*BC*_, and PD_*AB*_ were constructed based on the clustering of sensitivity profiles of levels of proteins B and C. For PD_*BC*_ and PD_*AB*_, we pick one and only one measurement candidate from each cluster; whereas for WD_*BC*_, all measurement candidates were chosen from the same cluster. PD_*BC*_, WD_*BC*_ and RD_*BC*_ denote designs where levels of proteins B and C are measured simultaneously, whereas PD_*AB*_ and RD_*AB*_ involve simultaneous measurement of levels of proteins A and B. The bottom panel displays the parameter estimation error corresponding to different realizations and its approximate distribution, for the different modalities. We consider 40 realizations of WD_*BC*_ and 100 realizations of each of the other modalities. The height of the bars depicts their mean. (b) The average estimation error of different sub-samplings of the 100 realizations of PD_*BC*_ and RD_*BC*_ is shown. (c) The distribution of different measurements represented in the 100 realizations of PD_*BC*_ and RD_*BC*_ is shown. The distributions, discretized in 12 bins, look similar have comparable entropy (PD_*BC*_: 3.55 bits; RD_*BC*_: 3.58 bits), and closely resemble a uniform distribution (3.59 bits).

We also observe that the performance of designs generated by PARSEC(k) depends on how closely certain design specifications are implemented. For instance, PD_*BC*_ optimized the measurements of proteins B and C based on their PSI. But if we quantify proteins A and B using the measurement time points optimized for the quantification of proteins B and C (PD_*AB*_), we observe a decrease in estimation accuracy compared to PD_*BC*_. However, PD_*AB*_ still outperforms RD_*AB*_ on average, with a 1.5 times lower error, which can be attributed to two factors: (a) PD_*BC*_ accounts for the sensitivity profiles of protein B, which is one of the variables measured in PD_*AB*_, and (b) a potential correlation between the sensitivity profiles of proteins A and C due to the coupled dynamics.

### PARSEC(k) tolerates parameter uncertainty

In the previous analysis, we optimized PD_*BC*_ using PSI evaluated at a guess of parameter values (GT_*G*_). But the guess might differ from the actual ground truth value (GT_*T*_). PARSEC(k) tolerates a significant disparity between the guess and actual value of ground truth but suffers when the guess is very bad as indicated by the accuracy ratio (Figure 5a), defined as the ratio of mean estimation error for random design to that for PARSEC(k) designs.

**Figure 5:**
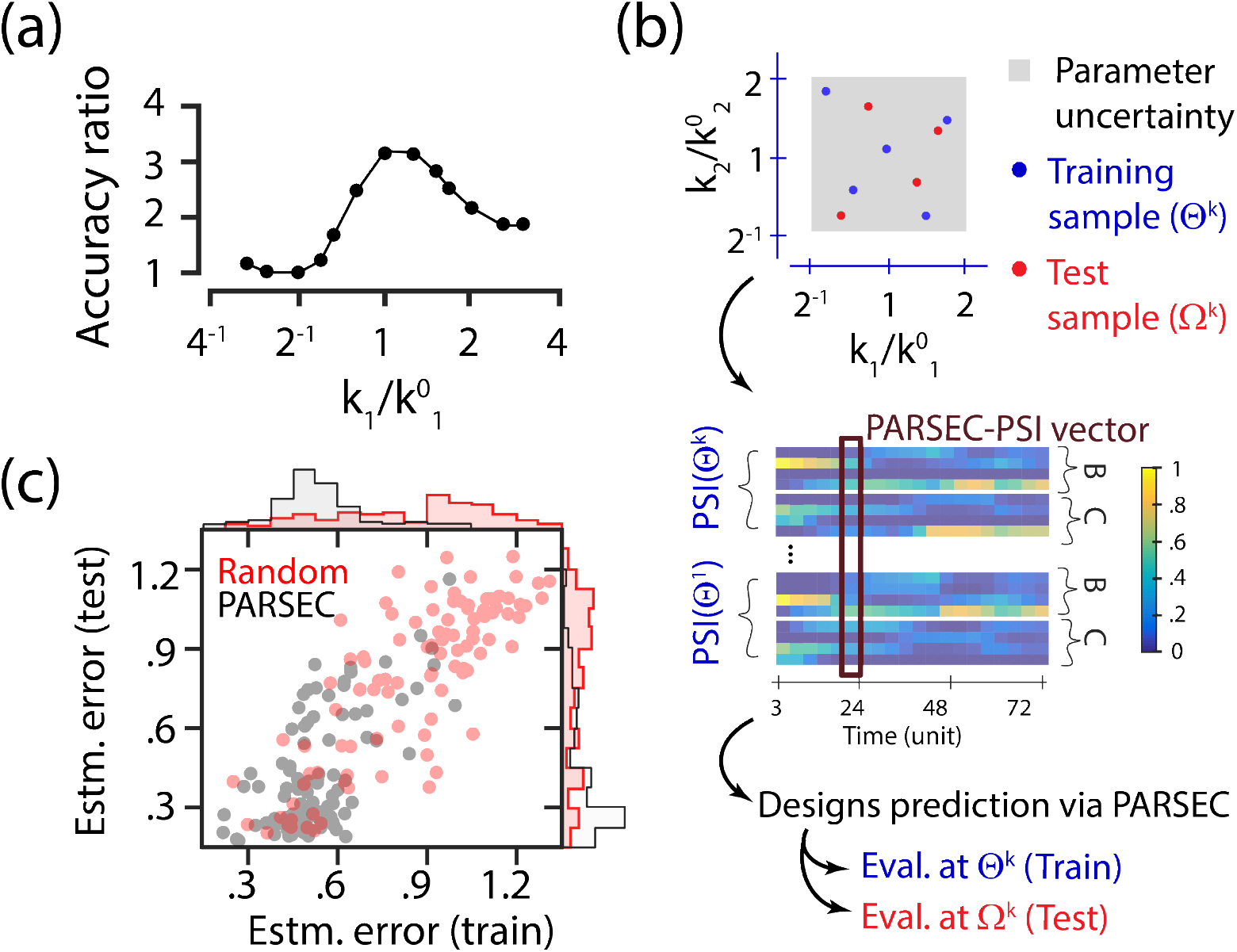
Robustness of PARSEC(k) designs. (a) Accuracy ratio evaluates how informative PARSEC(k) designs are compared to random designs. We see that when the ground truth guess (k^0^, used to implement PARSEC) is close to the actual ground truth (k_1_), PARSEC(k) designs lead to about 3.2 fold lower estimation error compared to random designs. This ratio (denoted as accuracy ratio) falls as k_1_ deviates from k^0^ (when the ground truth guess is bad). (b) However, PARSEC(k) can accommodate parameter uncertainty to identify generalist designs. It samples the range (Θ^*k*^) and clubs together the parameter sensitivity indices evaluated at these samples to create the PSI vector. The designs identified are based on the clustering of these PSI Vectors. The performance of PARSEC(k) is evaluated at Θ^*k*^’s (train analysis) and another set of parameter samples (Ω^*k*^, test analysis). (c) We use PARSEC(k) to accommodate a four-fold uncertainty in values of two of the six parameters characterizing the repressilator behavior and predict generalist or robust designs (schematically shown in Supplementary Figure S11). Using LHS, we identify five training samples (representing the uniform distribution in the log scales) and inform the PARSEC-PSI vectors; design evaluations are done at these training samples and four test samples. Train and test errors are calculated as the average estimation error across the training samples and test samples, respectively. The averaged estimation errors of 100 PARSEC(k) (black scatter) and random (red scatter) designs in train and test analyses are shown. The corresponding marginal is also shown. The plot highlights that PARSEC(k) designs are more informative. Interestingly, the PARSEC(k) designs that perform best in the training analysis also do so in the test analysis, thus identifying generalist designs.

Rather than informing designs using a single-point guess, PARSEC can do so for multiple guesses to accommodate uncertainty associated with parameter knowledge (Figure 5b). To do so, PARSEC samples the associated distribution for representative guesses called training samples (Θ^*k*^). PSI evaluated at the training samples are concatenated together to form the PARSEC-PSI vectors, used for clustering and subsequent design selection. Estimation errors for the PARSEC(k) and random designs are evaluated using data generated at the training samples (Θ^*k*^) and validated at another set of parameter samples (test samples, denoted as Ω^*k*^).

PARSEC(k) designs are on average twice as informative as random designs given the parameter uncertainty (Figure 5c). Interestingly, the top 5% of the PARSEC(k) designs in the training analysis, are also the best performers in the test analysis. Such designs robust to parameter uncertainty, can be good candidates for generalist designs. Moreover, comparable accuracy ratios for training and test samples (2 ± .85 and 2.05 ± .53, respectively) further emphasize the robustness of PARSEC(k) designs.

### Optimizing the sample size

In the previous analyses, we constrained the design search to those with pre-defined experiment sample size; here we optimize it. Random and PARSEC(k) designs lead to more accurate parameter estimation when their sample size increases (Figure 6a). However, the marginal gain in estimation accuracy decreases steadily (Supplementary Figure S12). Notably, PARSEC(k) designs of sample sizes five and higher exhibit similar performance on average (Figure 6a), suggesting that a five-measurement design would be both informative and economical.

**Figure 6:**
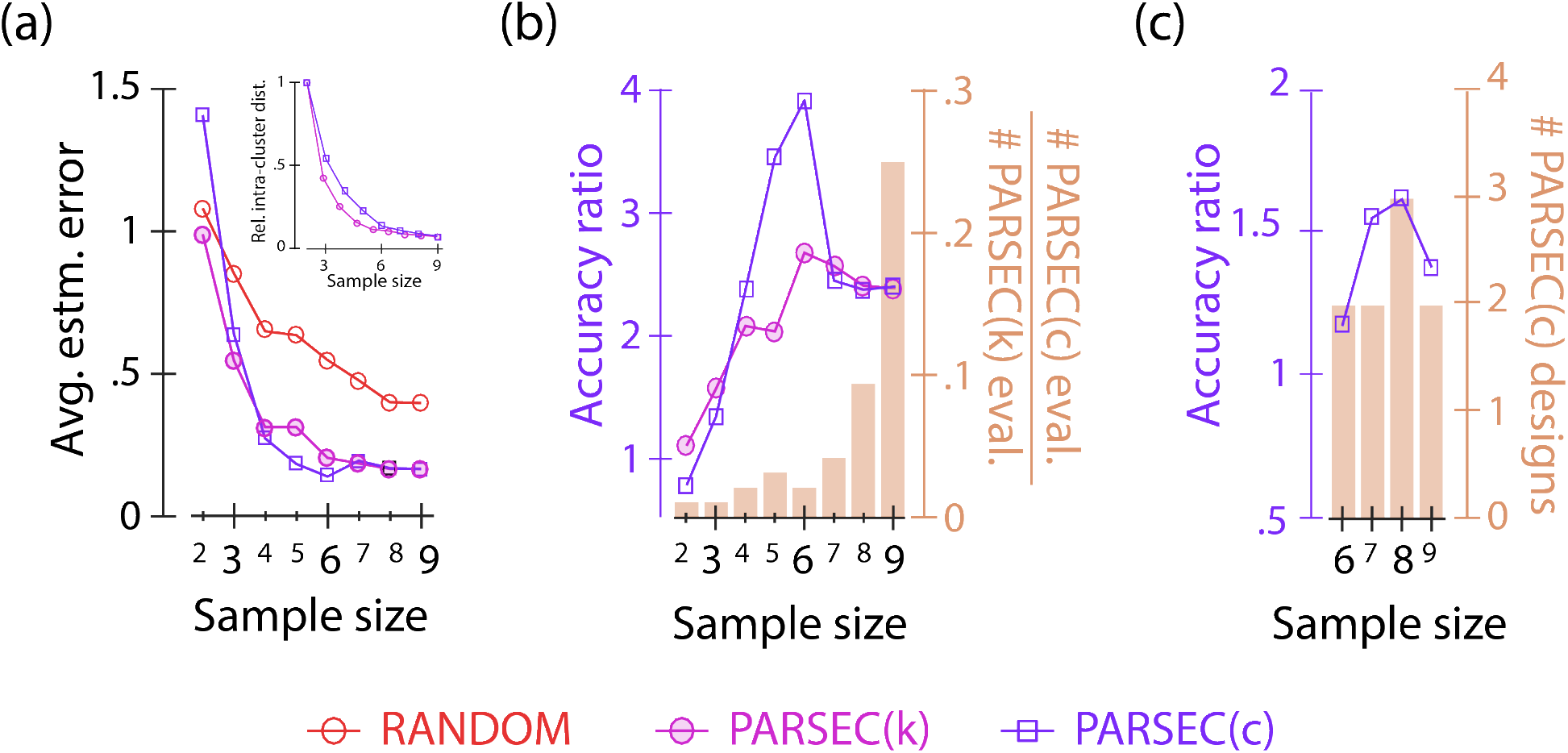
Optimizing experimental sample size. (a) We vary the sample size of the experiment and monitor the performance of PARSEC(k), PARSEC(c) and random designs, generalizing the design problem analyzed in Figure 4. The inset shows how the intra-cluster distance varies with number of clusters, due to k-means (open circle) and c-means (open square) clustering employed by PARSEC(k) and PARSEC(c) respectively. (b) We plot how the accuracy advantage of PARSEC(k) (open circle) and PARSEC(c) (open square), defined as the ratio of the mean estimation error of random designs and that of the corresponding PARSEC designs, vary with the number of clusters, which corresponds to the number of measurements. Here we construct 100 designs of each modality: random, PARSEC(k) and PARSEC(c). However, not all the 100 PARSEC(c) designs were unique resulting in a reduced downstream computation burden. Hence we also plot the ratio of PARSEC(c) designs and PARSEC(k) designs evaluated, as the fuzzy burden ratio.

As the number of clusters in PARSEC defines the experiment sample size, we posit that the clustering efficiency affects the information gain. PARSEC allows us to evaluate the relationship between the accuracy ratio and the clustering efficiency for a given sample size. In our example of designing experiments for the repressilator network, we find that the intra-cluster distance for the PARSEC-PSI vectors reaches an elbow point between five to six clusters, indicating optimal partitioning (Figure 6b). Interestingly, five clusters also correspond to repressilator PARSEC(k) designs with the highest accuracy ratio.

This suggests that clustering efficiency can serve as a valuable indicator for determining sample size for PARSEC, leading to high accuracy ratios. This arises due to two reasons. On one hand, using insufficient clusters limits the representation of PARSEC-PSI vectors in the PARSEC designs. Therefore, increasing the number of clusters significantly enhances the representation and information content of the designs, thereby improving the accuracy ratio. On the other hand, excessive clustering results in an over-representation of similar PARSEC-PSI vectors, resulting in only a modest increase in the information contents compared to that in random designs. Hence the advantage of using a PARSEC approach is likely to decrease with over-clustering. The observed correlation between clustering efficiency and accuracy ratio further validates the effectiveness of the PSI-clustering-based approach in experiment design.

While our implementation of PARSEC(k) provides a good estimate for optimal sample size, it still demands a substantial computational cost due to the evaluation of numerous designs. This cost escalates when considering parameter uncertainty across multiple parameter values. The multitude of predictions in PARSEC(k) arises from the discrete partitioning of continuous PARSEC-PSI vectors using the k-means clustering algorithm and subsequent candidate selection. An alternative approach is to employ the fuzzy c-means clustering (Supplementary Figure S13), as in PARSEC(c). This method results in fewer unique optima, drastically reducing the computational burden (Figures 6b). PARSEC(c) exhibits similar clustering convergence (Figure 6a: inset) and comparable, if not superior, estimation error and accuracy ratio profiles (Figures 6a and b) when varying sample sizes, all while requiring nearly twenty times fewer computations (Figure 6b). This enhanced efficiency renders sample size optimization for generalist design search manageable. We identified informative, generalist designs for various sample size experiments requiring a total of nine design evaluations (Figure 6c, Supplementary Figure S14), compared to the 100 designs evaluated to identify a generalist design of sample size six via PARSEC(k) (in Figure 5c).

## Methods

### Implementation of PARSEC

First, we identify the feasible measurement candidates according to design specifications and sample the parameter uncertainty selecting the ‘training samples’. PARSEC then evaluates the parameter sensitivity indices (PSI) for each variable within a measurement candidate across these training sample (step 1, Figure 1). These PSIs are concatenated into PARSEC-PSI vector for each candidate (Supplementary Figure S1). The candidates are grouped by similarity in their PARSEC-PSI vectors (step 2, Figure 1), guiding PARSEC’s final selection of measurement candidates (step 3, Figure 1). The candidates are selected such that each cluster is well-represented in the design.

We use Latin Hypercube Sampling (LHS [36]), a Monte-Carlo algorithm to efficiently sample multi-dimensional parameter spaces [37]. To generate N samples, LHS divides the distribution for each parameter into N equal probability intervals. Then it randomly picks an interval (without repetition) for each parameter and based on the distribution, chooses a random value from it. For sensitivity analysis, we use extended Fourier Amplitude Sensitivity Test (eFAST, [39, 40]). It is a variance-based method that gauges how parameter fluctuations contribute to uncertainty in the variable of interest. eFAST provides a global measure of sensitivity and captures non-linear relationships among parameters and outputs.

In our implementation, we apply either k-means or fuzzy c-means clustering algorithm. Suitable for partitioning continuous data into predetermined number of clusters, these unsupervised learning methods aim to minimize the total intra-cluster distance. k-means algorithm designates each data point to a single cluster. Whereas the c-means clustering algorithm assigns a probability of the data point belonging to each of the clusters. Upon c-means clustering, we guide the selection of measurement candidates using this probability measure of cluster membership. We iteratively select the measurement candidate corresponding to the highest degree of membership to any of the clusters (randomly breaking ties). Once selected, the measurement candidate and its corresponding cluster are ignored from the subsequent selection process. We denote to this flavor of PARSEC as PARSEC(c). Alternatively in PARSEC(k), we use k-means clustering algorithm and construct the design by randomly selecting a measurement candidate from each cluster. Properties such as silhouette scores can be used to bias selection of candidates from a cluster, alternate to the random analyzed in this study.

It’s important to note that the clustering and subsequent measurement selection steps in PARSEC can be stochastic. Thus we execute PARSEC to predict multiple designs, eventually selecting the most informative one. Designs are evaluated at one or more parameter values (step 4, Figure 1). At each parameter combination, we simulate the model and generate the data set according to the design being evaluated. We employ ABC-FAR to estimate the parameter values using the data set and calculate the estimation error.

### Implementation of ABC-FAR

ABC-FAR iteratively samples parameter combinations and rejects/accepts them based on associated χ^2^ values, which quantifies the deviation between data (R_*D*_) and corresponding model prediction 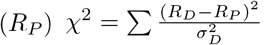, where the summation is over all data points and σ_*D*_ denotes the standard deviation in data R_*D*_.

ABC-FAR samples a noisy version of the current estimate of the distribution. However, the amplitude of noise added decreases with iteration (mimicking simulated annealing [38]). This improves exploration in early iterations without drastically slowing convergence in the later iterations. For sampling, we again rely on Latin Hypercube Sampling (LHS [36]). Typically each parameter is sampled independently, although correlations can be imposed using modified LHS algorithms [41, 42]. Independent sampling ignores the joint distribution of model parameters which can affect convergence. However, in an iteration, the algorithm allows us to include the combinations selected in the previous iteration during the parameter selection step (History-dependent update strategy, HDUS). This maintains the joint distribution to a certain extent, particularly when the current sampling of parameters is not as good as the previous selection. It also ensures that good combinations once sampled are not lost, guaranteeing a monotonic decrease in χ^2^ statistics with iteration. We can choose to not recall the previously selected parameter combinations (History independent update strategy, HIUS). HIUS saves on memory but does not guarantee convergence when weak rejection conditions (high FAR values) are used (Figures S6 and S7).

We also look at relationships among the components of the selected parameter combinations, like correlation and linear regression. This helps infer data-dependent properties like practical non-identifiability and associated degree of compensation between the parameters, respectively. Such measures capture linear and local relationships conditioned on values of other model parameters. Since parameter estimation and higher-order relationships are derived from a sampling-based approach, we infer their significance using a corresponding analysis of the dummy parameter. Although the dummy parameter doesn’t affect model prediction and hence χ^2^, we let ABC-FAR modify its distribution. Ideally, the distribution of the selected dummy parameter values should ideally mimic that of the sampled values. A deviation can help us identify computational artifacts and sampling biases. Higher-order relationships between a model parameter and the dummy parameter can help infer the statistical significance of relationships between model parameters.

### DoE for a three-gene repressilator system using PARSEC

A three-gene repressilator system (modeled by Elowitz and Leibler in 2000 [43]) involves the cyclic repression of gene expression, where one gene represses the expression of another. We use a simple ODE model that monitors the levels of the proteins expressed by these genes, with parameters representing the repression kinetic rates and thresholds. The protein levels show oscillatory dynamics.

In the analysis corresponding to Figure 4, we aim to identify a set of six synchronized measurements of protein B and C levels within 72 units of time (e.g. hours), for accurate parameter estimation. For this PARSEC calculates the parameter sensitivity indices (PSI) at a specific time point for levels of both proteins B and C. We concatenate the indices for proteins B and C to create a PARSEC-PSI vector for the measurement candidate at that time point. Based on the similarity of the PARSEC-PSI vectors, the measurement candidates are clustered via k-means clustering. We create six clusters, adhering to the constraint of having six measurements. From each cluster, we randomly select one representative candidate, which together forms the final design (PD_*BC*_). We predict 100 designs and compare them using ABC-FAR to identify the most informative one. Design comparison is based on the error in parameter estimation derived from the data emulating the design. To test the performance of PARSEC, we generate random designs (RD_*BC*_ and RD_*AB*_), non-specific designs (PD_*AB*_), and anti-PARSEC designs (WD_*BC*_). Each design constitutes six simultaneous measurements of the two variables (the protein levels). We generate a total of 100 designs for each mechanism, except for WD_*BC*_ where we generate only 40 designs.

Next, we study the robustness of PARSEC designs to uncertainty, characterized by a uniform distribution in log scale, over a four-fold range of values of two of the six model parameters (Figures 5a and b). PARSEC-PSI vector is constructed using PSI evaluated at five training values (Θ^*k*^) sampling the uncertainty via Latin Hypercube Sampling. Design evaluations are done at these five training samples and at another set of four test values (sampled via LHS). Since we consider a large dynamical range of uncertainty, we allow for a longer time frame of 120 units. Finally, while tuning the sample size (Figures 5c and d), we ignore uncertainty considerations evaluating PARSEC-PSI vectors at a single parameter guess and limit the duration of experiments to 72 units of time. Since the sample size is altered, we accordingly change the number of clusters which is used to partition the PARSEC-PSI vectors.

### Computational evaluation of a design

We evaluate design evaluation at specific values of model parameters. At the parameter combination, we simulate the model behavior based on the design specifications and sample this behavior according to the designs of interest to generate the associated data set. ABC-FAR is used to recover parameter values from the data. ABC-FAR selected parameter values are compared to the parameter combination used to generate the data, to calculate the estimation error associated with the design.

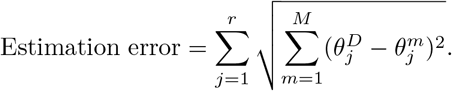

Here 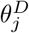 denotes the value of the j^*th*^ parameter of the combination used to generate data, 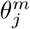 is the corresponding value of the m^*th*^ parameter combination selected by ABC-FAR. r and M indicate the number of free parameters to be estimated and the number of parameter combinations selected by ABC-FAR. In our implementation, ABC-FAR refines the posterior five times starting from a prior uniform distribution in log scale, spanning across two orders of magnitude. In each iteration, we sample 60,000 parameter combinations using a noisy version of the current estimate of marginal. We employ the History-dependent update strategy and a FAR value of 0.01. Furthermore, we add a uniform sampling noise 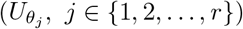 according to the following expression,

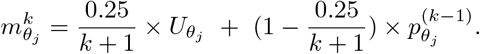

where 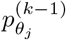 is the current estimate of marginal used to generate the sampling distribution 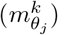, for the j^*th*^ free parameter (θ_*j*_) in the k^*th*^ iteration. Note 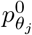 would be the prior or initial guess.

## Discussion

PARameter SEnsitivity Clustering-based algorithm (PARSEC) exploits the model structure and its parameter sensitivity to optimise the experimental design.First, even though the distribution of measurement candidates selected in PARSEC(k) is similar to that in the random designs (Figure 4c), the distribution of PARSEC-derived estimation errors is comparatively narrower and low (Figure 4a). This suggests that the degree of overlap between PARSEC-PSI vectors is more important to DoE than the individual characteristics of the measurements. Although we pick candidates randomly from each cluster, a systematic selection can further improve the efficiency of design search. Second, the parameter estimation error derived from designs decreases with a reduction in the overlap of the parameter sensitivity (PSI) vectors of measurements (comparison of PD_*BC*_ and WD_*BC*_, Figure 4a). Third, the efficiency of PSI clustering correlates with the accuracy ratio of the designs, as we vary the sample size (Figure 5d). This highlights the advantage of clustering the parameter sensitivity profiles in design generation and proposes a good guess for the optimal sample size for the experiment.

The critical ingredients of the algorithm are the PARSEC-PSI vectors. Our approach of vectorially conjoining the PSIs to create the PARSEC-PSI vectors allows us to identify generalist designs to mitigate parameter uncertainty. Here we concatenate the PARSEC-PSI evaluated at various parameter combinations (sampling the uncertainty), to construct the high-dimensional PARSEC-PSI vectors used for subsequent clustering and design selection. The robustness of designs to parameter uncertainty is a desirable property distinguishing PARSEC from locally optimal MBDoE approaches like Fisher’s information Matrix-based methods. Depending on the extent of uncertainty, PARSEC can either plan a ‘robust generalist’ design or a ‘locally optimized’ design. Nevertheless, robustness and accuracy are negatively affected as uncertainty increases (Supplementary Figure S11).

The vector-conjunction approach also allows us to easily implement various design constraints. For example, in our current analysis, we enforced simultaneous measurements of the different variables. Alternately, suppose that measurements of different variables have to be acquired at a fixed offset; then the PSI of the variables would be concatenated together in an appropriately staggered manner, to construct the PARSEC-PSI vectors. In case such restrictions are not there, one can treat each of the individual variables as different measurement candidates, during clustering and design selection (see Supplementary text SI S2 and Supplementary Figure S1). Furthermore the type of measurement also influences design specifications: certain variables may be more convenient/cheaper to measure than others. PARSEC can be implemented to identify variable combinations for economical and informative designs. Other specifications, like the choice of initial conditions, can also be tuned to improve designs for parameter estimation.

Optimizing the design parameters and design selection requires design evaluation. PARSEC employs Approximate Bayesian Computation-Fixed Acceptance Rate (ABC-FAR) algorithm for automated and fair comparison of designs. This is possible as ABC-FAR considers a global and relative sampling-rejection criterion (FAR), bypassing data-dependent stipulations. Additionally, a global parameter sampling method incorporating simulated annealing is used to improve the robustness of estimation and avoid local minima and biases. We have validated the algorithm for accuracy, reliability, efficiency, and robustness to initial bias and measurement noise. The efficiency can be further improved by adapting the FAR value for each iteration based on the improvement of convergence seen in the previous step. It has wide applicability and is amenable to the incorporation of available system-specific knowledge to improve convergence accuracy and speed (Supplementary Figure S10). Furthermore, we derive data-specific correlations and dependencies among the estimated parameter values, which can be used to infer properties, like practical identifiability. Like some of the other ABC methods, ABC-FAR can also be adapted for model selection by treating the identity of the model as a free parameter.

Parameter estimation-based design comparison, the final step in PARSEC is computationally intensive, much like other Bayesian Model-Based Design of Experiments (MBDoEs). However, the tight distribution of low estimation errors in PARSEC(k) indicates that its predicted designs are generally informative. This suggests that a limited number of design space samplings through PARSEC(k) should suffice, thereby reducing computational costs.

n alternative approach to mitigate computational burdens involves the use of fuzzy clustering. When considering the two variations of PARSEC presented here, it’s worth noting that PARSEC(c) designs (fuzzy clustering) represent a subset of PARSEC(k) designs. PARSEC(c) is highly efficient at identifying good designs but may overlook the most informative ones. In contrast, PARSEC(k) conducts a more exhaustive search, aiming to identify the most informative designs.

In summary, PARSEC leverages mathematical models and simulations to guide experimental design, maximizing the information obtained about model parameters within constraints. It can be applied to characterize complex systems or investigate the effects of perturbations. For instance, PARSEC can identify standardized, informative, and cost-effective drug screening experiments, facilitating the quantification and comparison of the impacts of different drugs on model parameters.

## Acknowledgement

We thank Dr. Mohit Kumar Jolly for sharing computational resources for carrying out the various analyses. We also appreciate the valuable feedback provided by Thiruvickraman Jothiprakasam, Ramya Boddepalli, Paras Jain, Sumanta Mukerjee and Suraj Jagtap on the manuscript.

## Author Contributions

Conceptualization, result interpretation and original draft preparation: HC and RR. Software and formal analysis: HC.

## Funding

This work was supported by the Indian Institute of Science Bangalore (RR), Wellcome Trust—DBT India Alliance intermediate fellowship (RR) and the Prime Minister Research Fellowship (HC).

## Competing interests

The authors declare that no competing interests exist.

## Supplementary Information

### SI S1 Constructing the PARSEC-PSI vector

Suppose *p*_*j*_ is the *j*^*th*^ of the ‘r’ model parameter being estimated and *V*_1_, *V*_2_, … *V*_*q*_ are the different variables whose levels can be measured. Parameter sensitivity is calculated only for the model parameters whose values are being estimated. To account for uncertainty we evaluate the PSI at multiple training samples (Θ^1^, Θ^2^, …, Θ^*k*^) representing the associated distribution. Now suppose we have a constraint where the variable V_*i*_ must be measured at a time point that is offset by Δ_*i*_ time units after the measurement of variable V_1_. So by definition Δ_1_ = 0. Furthermore, the case of simultaneous measurement of variables can be implemented by equating the offsets to zero, that is, Δ_2_ = Δ_3_ = … = Δ_*q*_ = 0.

Here we demonstrate how to construct the PARSEC-PSI vector for the general offset situation (top panel, SFigure S1). Here a measurement candidate is defined as the sequence of measurement of the q variables at their proper offset. Let the measurement time point be labeled using, t, the time point where V_1_ is measured. We have a PARSEC-PSI vector for each of the measurement time points. Parsing the top panel of Sfig from right: SI_*p*_(V_*k*_(t + Δ_*k*_)) |_Θ*m*_ denotes the parameter sensitivity index of V_*k*_ at time t + Δ_*k*_ towards the p^*th*^ model parameters evaluated at Θ^*m*^. These indices form the components of PSI[V_*k*_(t + Δ_*k*_)] |_Θ*m*_ representing the parameter sensitivity of the variable V_*k*_ at time t + Δ_*k*_ evaluated at parameter value Θ^*m*^. PSI (V_*k*_(t)) |_Θ*m*_ is a r-dimensional vector. These vectors for different variables accounting for appropriate offsets are chained together to give PSI(t, Θ^*m*^), the parameter sensitivity profile of the entire measurement candidate evaluated at Θ^*m*^. Finally, PSI(t, Θ^*m*^) for all the training samples (m = 1, 2, …, n) are conjoined to form the PARSEC-PSI(t), which will be used for clustering. The PARSEC-PSI vector is an (r.q.n)-dimensional vector, accounting for the ‘r’ free parameters, the ‘q’ variables constituting a measurement, and ‘n’ training samples that accommodate the uncertainty.

**Figure S1:**
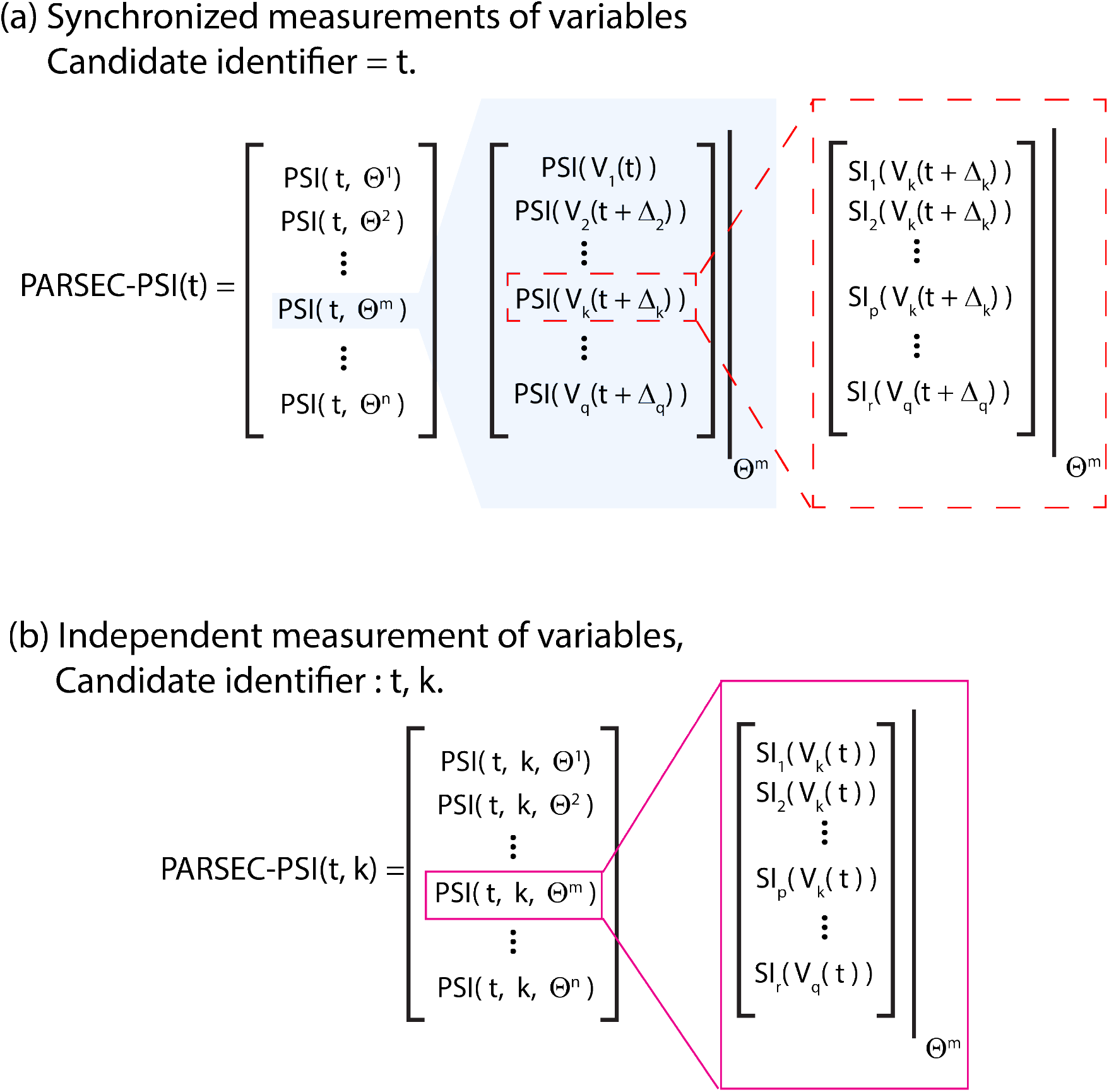
Constructing the PARSEC-PSI vector. PARSEC relies on the clustering of the PARSEC-PSI vectors. Thus these vectors are the main functional components of the PARSEC algorithm. PARSEC allows us to design these vectors to account for parameter uncertainty and various experiment design constraints. For example, suppose we want to characterize a kinetic system by measuring V_1_, V_2_, … V_*q*_. We show the procedure for constructing the PARSEC-PSI vectors for two scenarios: (top panel) when designs must consider synchronized measurement of all the variables, then the only measurements are defined by the time of measurement of one of the variables, and (bottom panel) when the measurements of one variable do not depend on the measurement of the other variables, then the designs are more flexible; the measurements are defined using the variable label and the time label (bottom panel).

Alternatively, if the constraint of offset and collective measurement of the variable is relaxed, the construction of the PARSEC-PSI vectors is simpler (SFigure S1). In this flexible scheme, if and when a variable is measured is not constrained by the measurement of the others. Thus, a measurement candidate is characterized by which variable is being considered (variable label = k) and the time point of measurement (time label = t). Thus, the corresponding PARSEC-PSI vector, PARSEC-PSI(t, k), is a concatenation of ‘n’ PSI vectors - PSI(t, k, Θ^1^), PSI(t, k, Θ^2^), …, PSI(t, k, Θ^*n*^) - where PSI(t, k, Θ^*m*^) is the parameter sensitivity vector for the variable V_*k*_ at time t evaluated at training sample Θ^*m*^ (for m = 1, 2, … n). PSI(t, k, Θ^*m*^) is a ‘r’ dimensional vector, whose p^*th*^ component is SI_*p*_(V_*k*_) |_Θ*m*_ denoting the parameter sensitivity index of V_*k*_ at time t towards the p^*th*^ model parameters evaluated at Θ^*m*^. So each of the PARSEC-PSI vectors, PARSEC-PSI(t, k), is an (r.n)-dimensional vector, accounting for the ‘r’ free parameters and ‘n’ training samples that accommodate the uncertainty.

### SI S2 ABC-FAR algorithm for model parameter estimation

We propose a likelihood-free, Approximate Bayesian computation-(ABC) based algorithm for fitting model and estimating corresponding parameter values (Figure 2a, Main text). We use Monte-Carlo sampling technique to avoid explicitly calculating the (typically intractable) likelihood as a function of parameter value. Instead we calculate the deviation between model prediction and data (denoted as χ^2^), by simulating the model for sampled parameter combinations. We select a fixed fraction of the number of the sampled parameter combinations with the lowest χ^2^ value to update the current distribution. We denote the value of this fixed fraction as the Fixed Acceptance Rate or FAR. The process of sampling from the current distribution, selection of parameter combination and updating the current distribution, is repeated to narrow down the plausible parameter value range. This concentrates computations in the later iterations to narrower ranges where the parameter values are more likely to lie, making the algorithm efficient. In our analysis, we include an additional dummy parameter which doesn’t affect model prediction but undergoes the same algorithmic treatment as the model parameters being estimated. It is used to (a) detect artefacts, if any introduced by the algorithm, and (b) inform on significance of properties of estimated model parameters.

Here we describe the parameter sampling and acceptance modules of the algorithm. Suppose the model being fit, has k parameters. In each iteration, N realizations of parameter combination is sampled using the current estimate of the marginal of the k model parameters. For the first iteration, the current estimate of marginal would be the prior distribution (initial guess), whereas in subsequent iterations, the current estimate is the posterior estimated in the previous iteration. The algorithm employs Latin Hyper-Cube Sampling (LHS, [1]) to effectively scan the (potentially high-dimensional) parameter space. A noise, whose magnitude decreases with iteration, is added to the distribution each time before sampling. This helps in exploring the parameter space better in the earlier iterations without hindering the convergence, significantly, in the later ones, like in simulated annealing [2]. Adding a sampling noise helps avoid local minima in the earlier iterations and improve robustness of the estimation.

The model is simulated for each of the N parameter combinations sampled. Appropriate model predictions are compared to the data, to estimate χ^2^ statistics as a measure of the deviation. χ^2^ is the weighted sum of squared of difference between prediction and data, weighted by variance in measurement. Desirable model fits should closely match the data. We select the M (= N × FAR) parameter combinations corresponding to the lowest χ^2^ (best agreement between prediction and data) to update the estimate of distribution of the parameter values.

From the second iteration and onwards, we can choose to include previously selected parameter combinations, while sorting the N sampled combinations and updating the posterior in an iteration. This ‘history-dependent update strategy’ (HDUS) ensures that newly sampled parameter combinations are only accepted if they are better than those selected earlier. On the other hand, the ‘history-independent update strategy’ (HIUS) selects parameter combinations solely from the current sampling to update the posterior. Irrespective of the update strategy, in each iteration, N model parameter combinations are sampled and only M parameter combinations are selected. This maintains an acceptance rate of FAR = M/N, in each iteration. However the total number of sampling and thus computations increases every iteration. Thus the computation efficiency falls inversely with iteration and is characterized by the effective acceptance rate 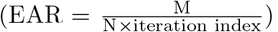. In this article, we discuss how the choice of update strategy and FAR values affect efficiency and convergence of the algorithm, after we demonstrate the working of the algorithm.

Keeping the initial condition fixed, we employ the ABC-FAR algorithm, to fit the model to the data. We consider N = 10^4^ samplings and a FAR (fixed acceptance rate) of 2.5% to update the posterior, which is iterated eight times. We use the. Thus second iteration and onwards, the algorithm selects 250 (M = N×FAR = 250) best parameter combinations from a collection of ‘N (sampled in the current iteration) + M (selected in the previous iteration)’ combinations.

### SI S3 Description of the systems considered

#### Prey-predator system (Lotka-Votka model)

We look at the popular Lotka Volterra model, which is typically used to benchmark parameter estimation algorithms (as in [3]). The model tries to explain the population dynamics due to the interaction between prey and predator species. In the model, the population of prey (N) increase exponentially in absence of the predators (associated parameter: a). If predators are present, they interact with and consume the prey. This leads to a decrease in the prey population (associated parameter: b), but promotes the proliferation of predators (associated parameter: c). However, in absence of preys, the predator population (P) decays exponentially (associated parameter: δ), due to lack of food/resources.

**Lotka-Votka model**

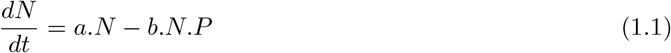

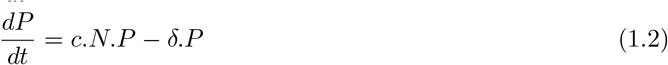

#### Three-gene repressilator system

The system consists of three genes regulating each other’s expression. Expression of one gene produces a protein, that represses the expression of the other gene in cyclic manner.

**GRN model**

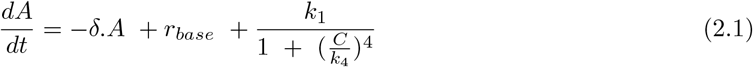

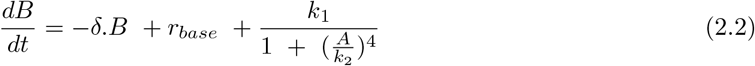

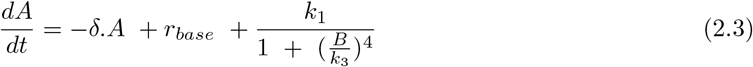

#### Coherent feed forward (type-1) loop - FFL-1

We look at a coherent feed-forward (type-1) network for gene regulation. Molecules of species S (population size - S) activates the production of protein A (population size - A). Presence of A induces the degradation of S. Protein A activates the production of a protein B (population size - B). Presence of both the proteins, A and B, promotes the production of another protein C (population size - C). Additionally we also consider a natural decay/dilution kinetics for the proteins A, B and C (rate parameter - δ).

**FFL-1 model**

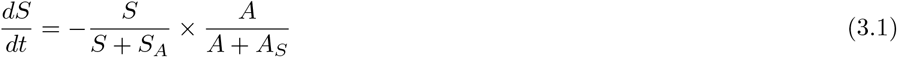

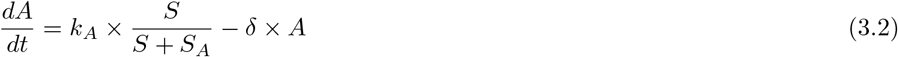

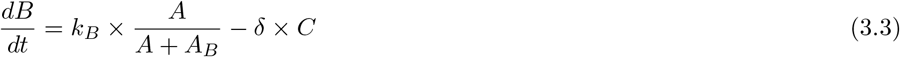

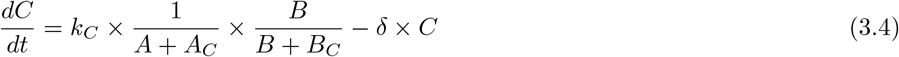

### SI S4 Data generation for fitting

Data used for fitting were computationally generated by simulating the model (on MATLAB). We assume values for the model parameters (Ground truth value, GT) and initial conditions, and simulate the model. The relevant model predictions sampled at the time points of interest constitute the data.

To emulate measurement error, we pick data points from a distribution centered around the model prediction. For example, suppose the model prediction for variable, V at time t, is V (t) = V_1_. A corresponding noisy data would be V_*N*_ (t) = V_2_ Normal[V_1_, σ], where Normal[a, b] is a normal distribution with mean a and variance b^2^.

We consider two dynamical (biological) systems to demonstrate various aspects of ABC-FAR, the parameter estimation algorithm developed. We consider (a) a two-species prey-predator system, and (b) a coherent feed forward (type-1) loop, described in the previous section.

**Table S1:** Bounds used during fitting and the GT values used for data generation.

| Parameter | Lower bound | Lower bound | GT<sup>#</sup> |
| --- | --- | --- | --- |
| **Lotka-Volterra model** |  |  |  |
| $\log_{10}(a)$ | -1 | 1 | $\log_{10}(1)$ |
| $\log_{10}(b)$ | -2.5 | 0 | $\log_{10}(0.02)$ |
| $\log_{10}(c)$ | -1 | 1 | $\log_{10}(0.25)$ |
| $\log_{10}(\delta)$ | -2 | 0 | $\log_{10}(0.3)$ |
| dummy | 0 | 1 |  |
| **FFL-1 model** |  |  |  |
| $\log_{10}(k_A)$ | -1 | 2 | $\log_{10}(2.5)$ |
| $\log_{10}(S_A)$ | -1 | 2 | $\log_{10}(3)$ |
| $\log_{10}(A_S)$ | -1 | 2 | $\log_{10}(5)$ |
| $\log_{10}(\delta)$ | -2 | 1 | $\log_{10}(0.1)$ |
| $\log_{10}(k_B)$ | -1 | 2 | $\log_{10}(2.5)$ |
| $\log_{10}(k_C)$ | -1 | 2 | $\log_{10}(5)$ |
| $\log_{10}(A_B)$ | -1 | 2 | $\log_{10}(15)$ |
| $\log_{10}(A_C)$ | -1 | 2 | $\log_{10}(2)$ |
| $\log_{10}(B_C)$ | -1 | 2 | $\log_{10}(3)$ |
| dummy | -1 | 2 |  |
| **GRN model** |  |  |  |
| $\delta$ | | | 0.1 |
| $r_{base}$ | | | 0.05 |
| $k_1$ | | | 1 |
| $k_2$ | | | 2 |
| $k_3$ | | | 3 |
| $k_4$ | | | 4 |

**Table S2:** Simulation conditions to generate synthetic data.

|  |  |
| --- | --- |
| **Lotka-Volterra model** |  |
| Simulation time | 0 - 30 |
| Initial conditions | Prey population size: 75 |
|  | Predator population size: 20 |
| **FFL-1 model** |  |
| Simulation time | 0 - 150 |
| Initial conditions | We start with all model variables set to zero except the level of S which was set to 20. |

For description of the model parameters refer to the discussion on corresponding models. GT^#^ refers to the ground truth values which was used to generate synthetic data.

### SI S5 Correlation-based statistics corresponding to selected parameter values

We posit that PnI(p, q) captures practical non-identifiability between the p^*th*^ and the q^*th*^ model parameters. The presence of correlation would indicate that changes in predictions of interest (in the context of data) due to perturbations in the p^*th*^ parameter can be compensated via appropriate changes in the q^*th*^ parameter, and vice-versa. For example, if PnI(p, q) is positive, then perturbation to predictions of interest due a decrease in value of p^*th*^ parameter would be buffered by a decrease in value of q^*th*^ parameter. Since this is Bayesian-based measure, we call it Bayesian Practical Identifiability. Note that the correlations would primarily capture local and linear effects.

Additionally we can also look at correlation between selected parameter values and corresponding χ^2^ values. Let ConvCorr(p) evaluates the correlation between the value of the p^*th*^ parameter in the selected combination and the corresponding χ^2^ value. A positive (negative) correlation suggests that among the selected values of the parameter, the ones with lower (higher) value correspond to predictions with higher (lower) deviation from data. Thus the presence of correlation indicate that further refinement in (the centrality measure of) the posterior is needed. Note that the correlation between χ^2^ and a model parameter is a local property, conditional on the other model parameter values being fixed. We can look at a more global measure - TCC. TCC denotes the total magnitude of correlation between selected parameter values and χ^2^ summed over all model parameter. If TCC decays to zero, then each term in the sum goes to zero, suggesting convergence to a minima for each of the parameters. Simultaneous local convergence in turn implies global convergence in (the centrality measure of) posterior estimation. Thus we posit the decay of TCC to zero to indicate convergence of model fitting. Since TCC based convergence is Bayesian computation based, we refer to it as a Bayesian correlation-convergence criterion.

We borrow the same notations as used in the main text. 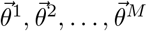 are the M parameter combinations selected based on their χ^2^ values. Let 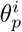 denote the value of the p^*th*^ model parameter in the i^*th*^ combination. Further let 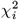 denote the χ^2^ statistics corresponding to the i^*th*^ parameter combination, 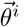. The correlative measures are evaluated as:

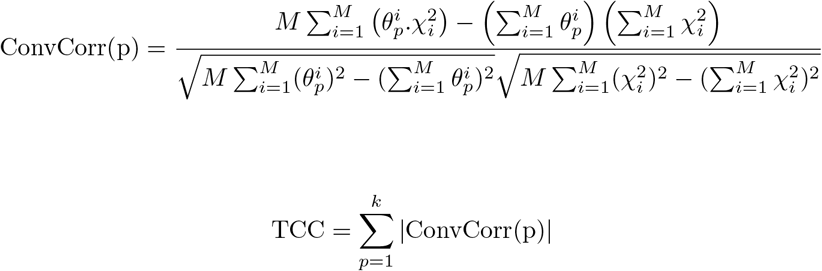

### SI S6 Efficiency of parameter estimation algorithm

We evaluate (a) minimum/maximum χ^2^ statistics, and (b) Effective Acceptance Rate (EAR), to fairly compare the performance of different schema of ABC-FAR with each other and with the reported performance of ABC-SMC. Minimum and maximum χ^2^ statistics looks at the minimum and maximum of the χ^2^ values corresponding to the parameter combinations selected at every iteration of the estimation procedure. Central tendencies, like mean or median, of the (selected) χ^2^ values can also be used. These χ^2^ statistics indicate the goodness of fit, or how well the model predictions mimic the data used for fitting. On the other hand, the Effective Acceptance Rate (EAR) looks at the (inverse of the) cumulative computational complexity. EAR is proportional to the total number of parameter combinations evaluated or the total number of times the model was simulated.

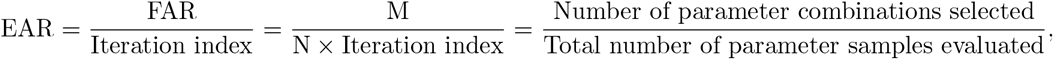

where N and M are the numbers of parameter combinations sampled and selected in an iteration respectively.

An algorithm or scheme leads to a good/accurate fit if it can identify parameter combinations corresponding to low χ^2^ values. Comparing schemes that reach a particular threshold of χ^2^ statistics, the one with higher EAR (that is, less computational cost) is considered more efficient than the other. Comparing the different schemes of our algorithm was straightforward as we use the same χ^2^ function and identical value of M (the number of parameter combinations selected in an iteration). However, we have to normalize for rejection criterion and differences in the χ^2^ function while comparing different algorithms. The ABC-SMC study reports their performance for a single iteration implementation along with the performance of an optimized realization of their algorithmic parameters. We consider the following normalization,

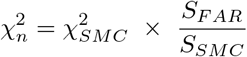

where 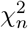 is the normalization of 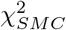 (from ABC-SMC paper) based on S_*F AR*_ and S_*SMC*_ which are the χ^2^ statistics corresponding to single iteration implementation of ABC-FAR and ABC-SMC algorithms respectively. To normalize for different rejection criteria, we calculate S_*F AR*_ imposing a FAR value of 7 × 10^*−*4^, so that the EAR corresponding to both S_*F AR*_ and S_*SMC*_ are same.

### SI S7 Compatibility of ABC-FAR to utilize system-structure to improve convergence

In the previous section we saw how algorithm-based strategies affect efficiency of parameter estimation. In some situations, we can also utilize system-specific knowledge to speed up and/or improve convergence, for example, when we have partial knowledge about parameter value distribution or the mathematical structure of the model allows us to define the fitting problem cleverly. We demonstrate the compatibility of our program to exploit such information to speed up and improve convergence. We demonstrate how we can do so for the FFL1 system, described in the previous section.

According to the model, the dynamics of levels of S and A are influenced only by each other. Thus we can focus on the sub-model and data associated with only their dynamics to estimate the associated parameters - log_10_(k_*A*_), log_10_(S_*A*_), log_10_(A_*S*_) and log_10_(δ)). We label this sub-problem as ‘H1’, where we ignore the parts of the model associated with dynamics of B and C. Next we observe that dynamics of B is affected by levels of S and A, but not by the level of C. Thus we define sub-problem ‘H2’, where we augment the sub-problem H1 with the dynamics of B and the additional parameters associated with it - log_10_(k_*B*_) and log_10_(A_*B*_)). The posterior estimated in H1, for the first four parameters can be used as prior in H2. After H2, we finally consider the entire model in sub-problem ‘H3’. For all the older parameters, we use the posterior estimated for the parameters in H2, as the prior for respective parameters in H2. We denote this strategy of defining sub-problems as Hierarchical calibration (hCal). We compare its efficiency against the one-shot calibration (oCal) strategy where we fit the entire model at a single go.

In this analysis, parameter values in log scale to explore a large dynamical range. Also, whenever a parameter is estimated for the first time in either of the calibration strategies, we use a uniform prior. So for hCal, we use a uniform prior for (a) log_10_(k_*A*_), log_10_(S_*A*_), log_10_(A_*S*_) and log_10_(δ)) in H1, (b) log_10_(k_*B*_) and log_10_(A_*B*_)) in H2, (c) log_10_(k_*C*_), log_10_(A_*C*_)) and log_10_(B_*C*_) in H3; we use uniform priors for all the parameters in the oCal strategy. We use two iterations each for H1 and H2, four for H3, and eight for oCal; thus a fair comparison can only be made after the fifth iteration when the entire model is considered and χ^2^ functions used in both the strategies are identical. Other algorithm options are kept identical: FAR = 10%, HDUS, parallel execution. To compare efficiency we look at the cumulative execution of time (cTime) and χ^2^ values for selected parameter combinations.

We see that both the calibration strategies fits the model to data (SFigure S7), but hCal is more efficient and accurate compared to oCal. For example, fifth iteration onwards (a) the χ^2^ values corresponding to the selected parameters, and (b) the cumulative execution time, are lower for hCal compared to oCal. Actually, hCal outperforms oCal by almost an iteration in both these measures. cTime corresponding to the seventh iteration of hCal compares with that of the sixth iteration of oCal, whereas χ^2^ distribution corresponding to the seventh iteration of hCal is as good as that of the eighth iteration of oCal (SFigure S7). The better performance of hCal can be attributed to (a) fewer calculations required for simulation and χ^2^ evaluation in H1 and H2, (b) lower dimensionality of parameter space for H1 and H2, and (c) propagation of information about parameters from one sub-problem to next.

Our algorithm was conveniently adapted and easily set-up to execute the hCal strategy. We didn’t have to define different sets of parameter rejection threshold for the three sub-problems associated with hCal, a convenient advantage over the typically used ABC-based parameter estimation algorithms.

## Supplementary figures

**Figure S2:**
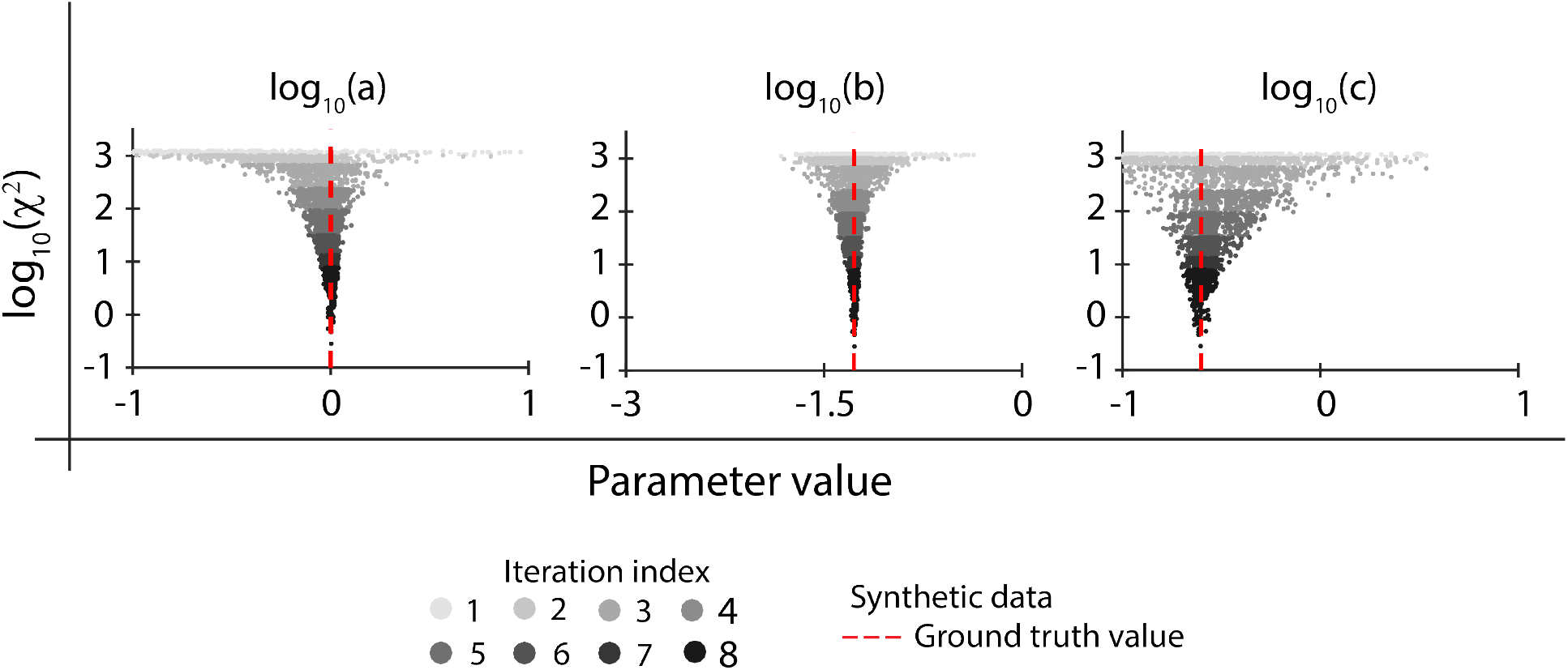
Scatter-plot showing the the parameter values selected and the corresponding χ^2^ value, in each iteration of fitting the Lotka-Volterra model. The values of iteration index are represented by shades of gray. Here the prey-predator model is fit to the synthetic data, generated using the parameter values denoted as ‘Ground truth’ values, depicted by the red dashed line.

**Figure S3:**
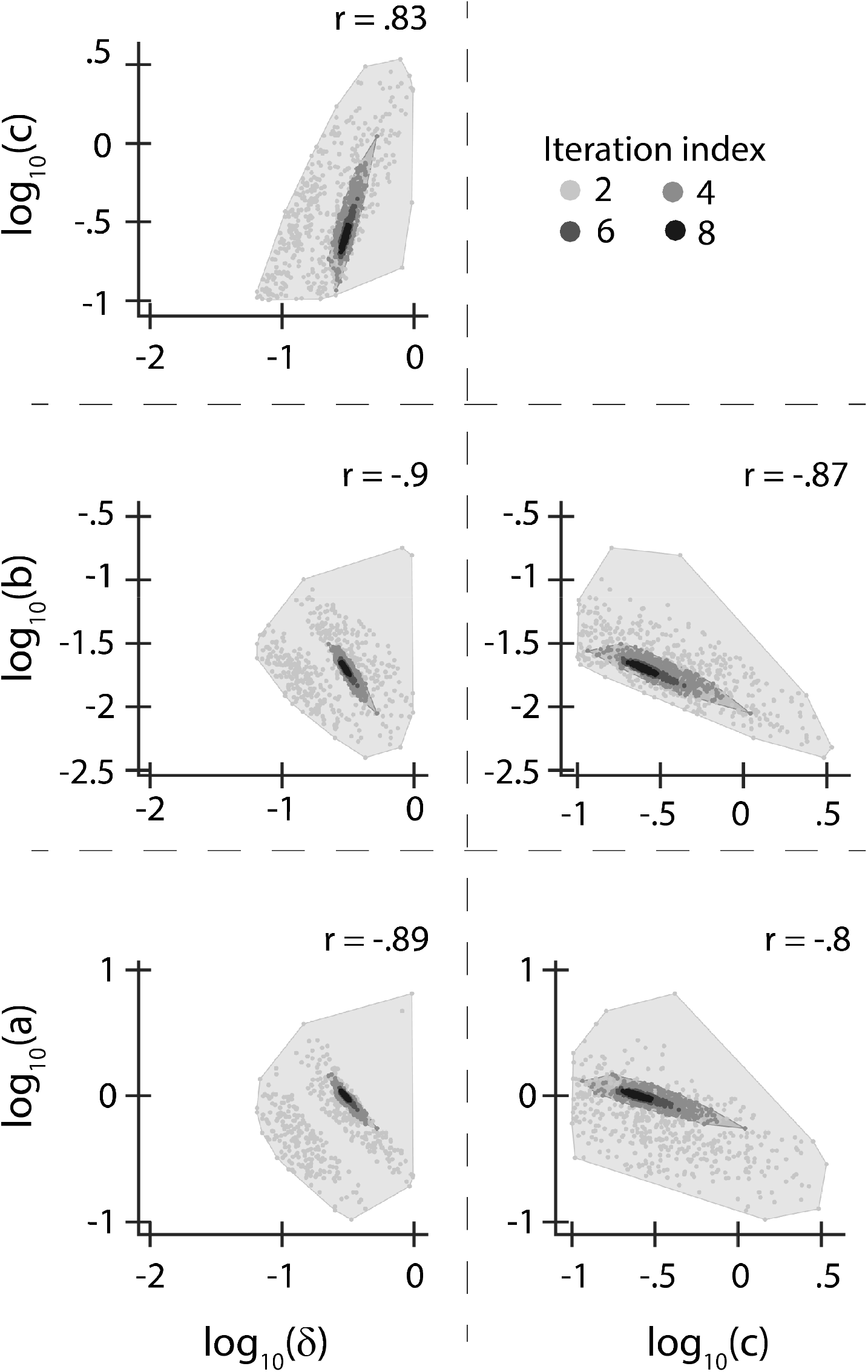
2D scatter-plots (and the convex hull boundary) of parameter combinations selected in the second, fourth, sixth and eighth iterations of fitting the Lotka-Volterra model. The iterations are represented by the shades of gray. r-value shown in the top-right corner for each sub-figure shows the correlation between the values, of corresponding parameter-pair, selected in the final (eighth) iteration.

**Figure S4:**
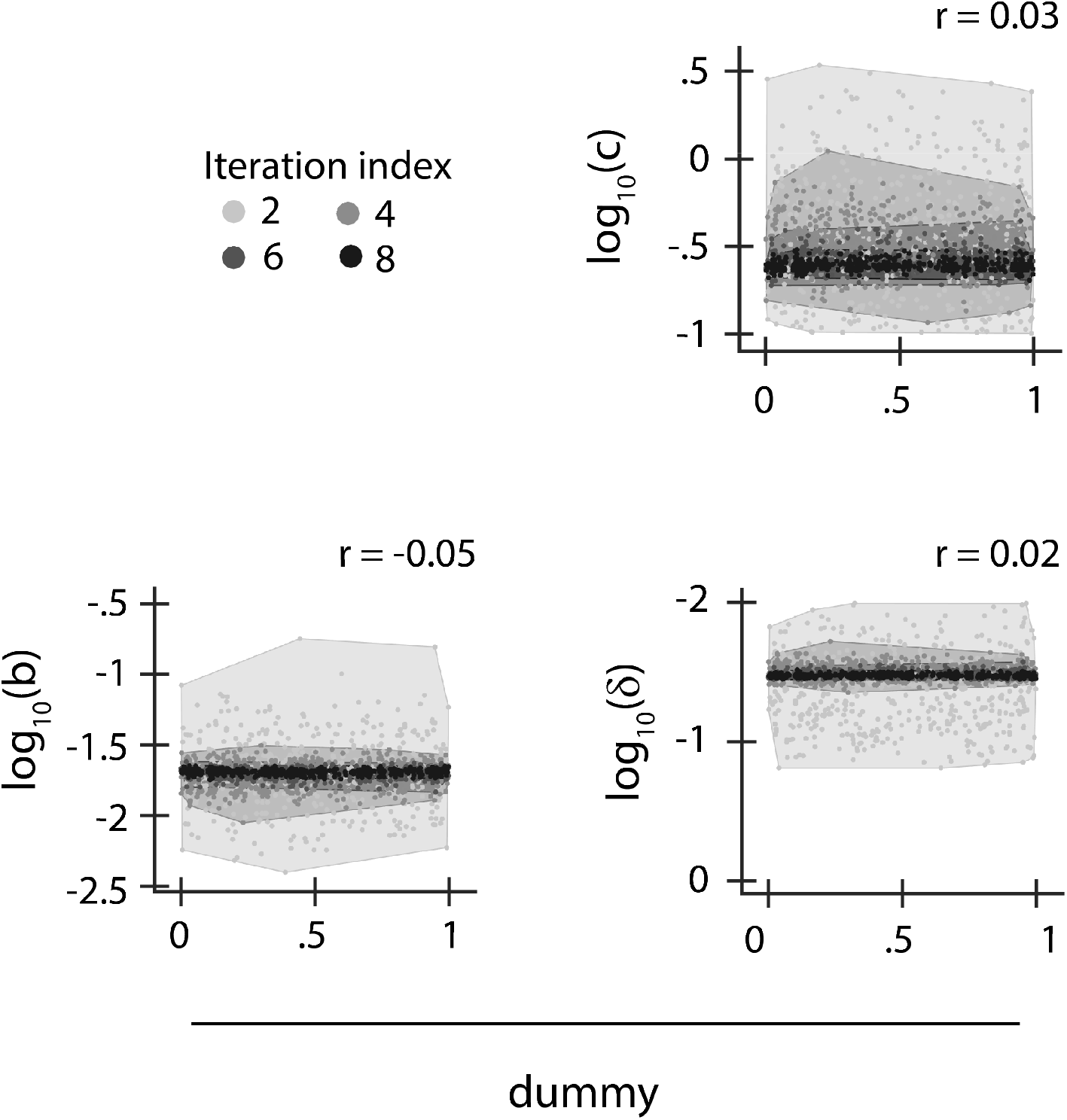
2D scatter-plots (and the convex hull boundary) of parameter combinations selected in the second, fourth, sixth and eighth iterations of fitting the Lotka-Volterra model. The iterations are represented by the shades of gray. r-value shown in the top-right corner for each sub-figure shows the correlation between the value of corresponding parameter and dummy parameter value selected in the final (eighth) iteration.

**Figure S5:**
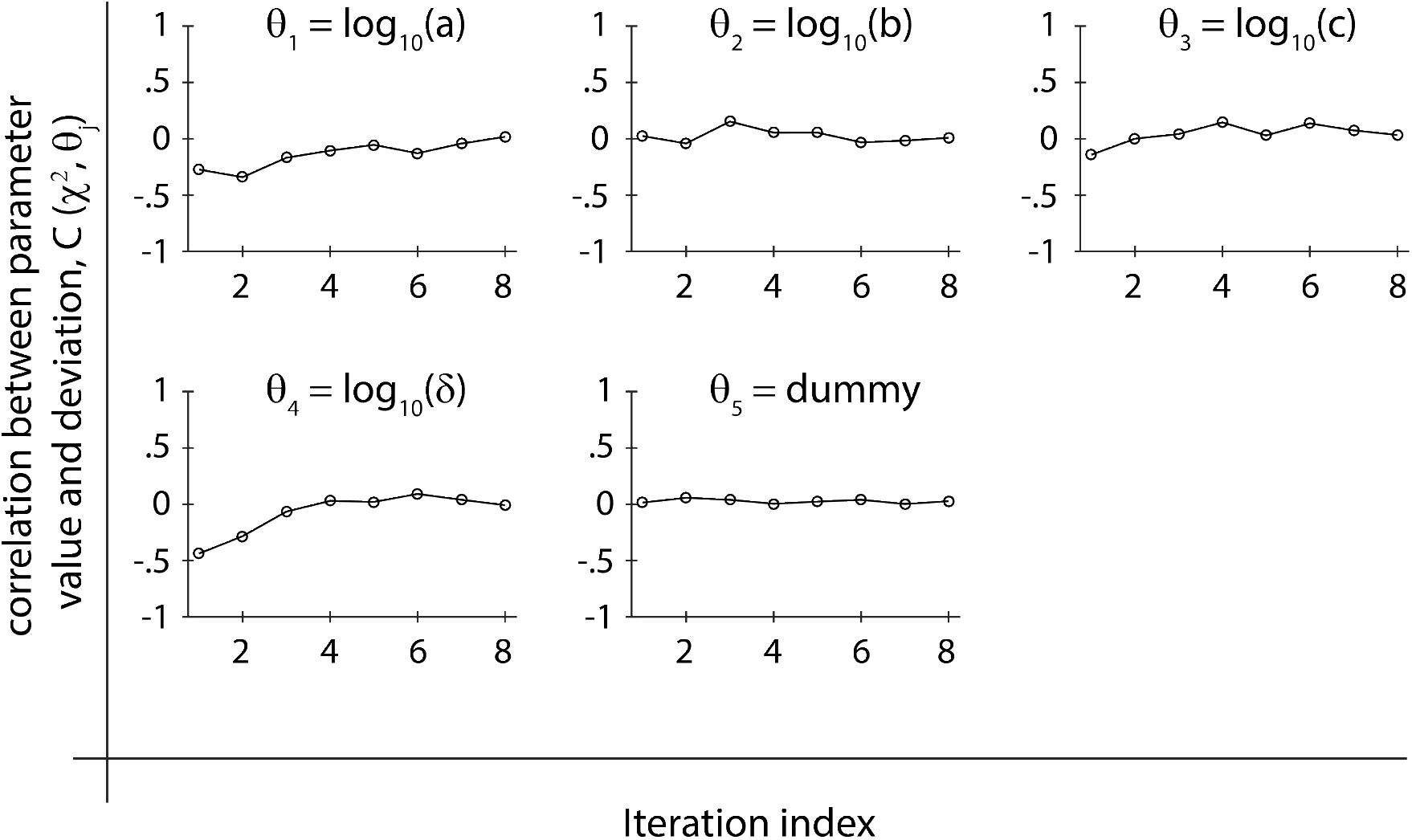
Variation of the correlation between the χ_2_ and the values of the parameters, selected after every iteration, while fitting the Lotka-Volterra model.

**Figure S6:**
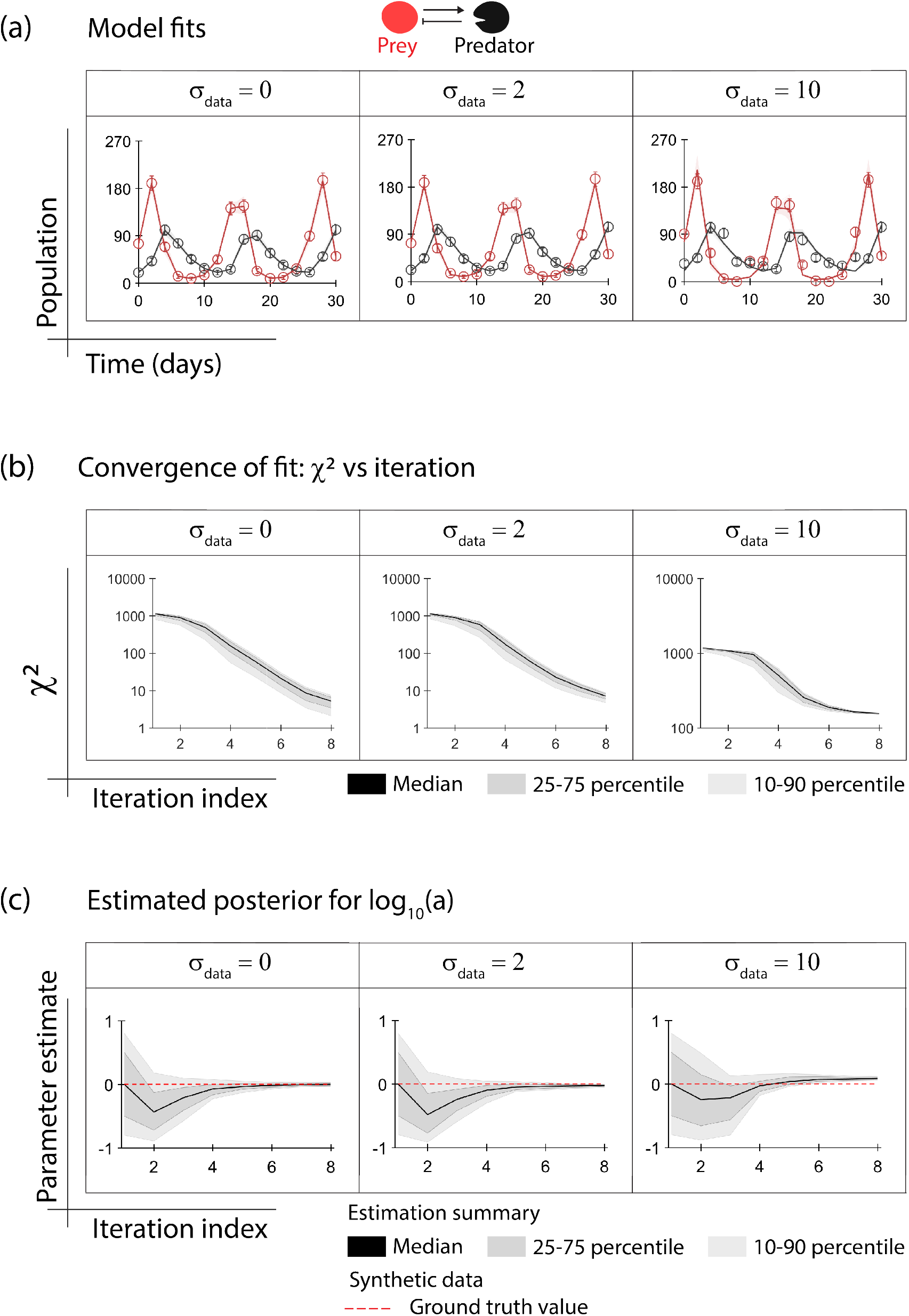
Model fitting and parameter estimation using noisy data. Data used for fitting the Lotka-Volterra model was augmented with noise to mimic measurement error (σ_data_) in measuring population size. (a) The model predictions due to parameter combinations selected at the final iteration (thin lines) explain the noisy data (open circle) well. The thick lines show the mean of predictions due to the selected parameter combinations. (b) Statistics for the distribution of χ^2^ corresponding to the parameter combinations selected at each iteration. (c) Statistics for the posterior of log_10_(a) estimated at each iteration.

**Figure S7:**
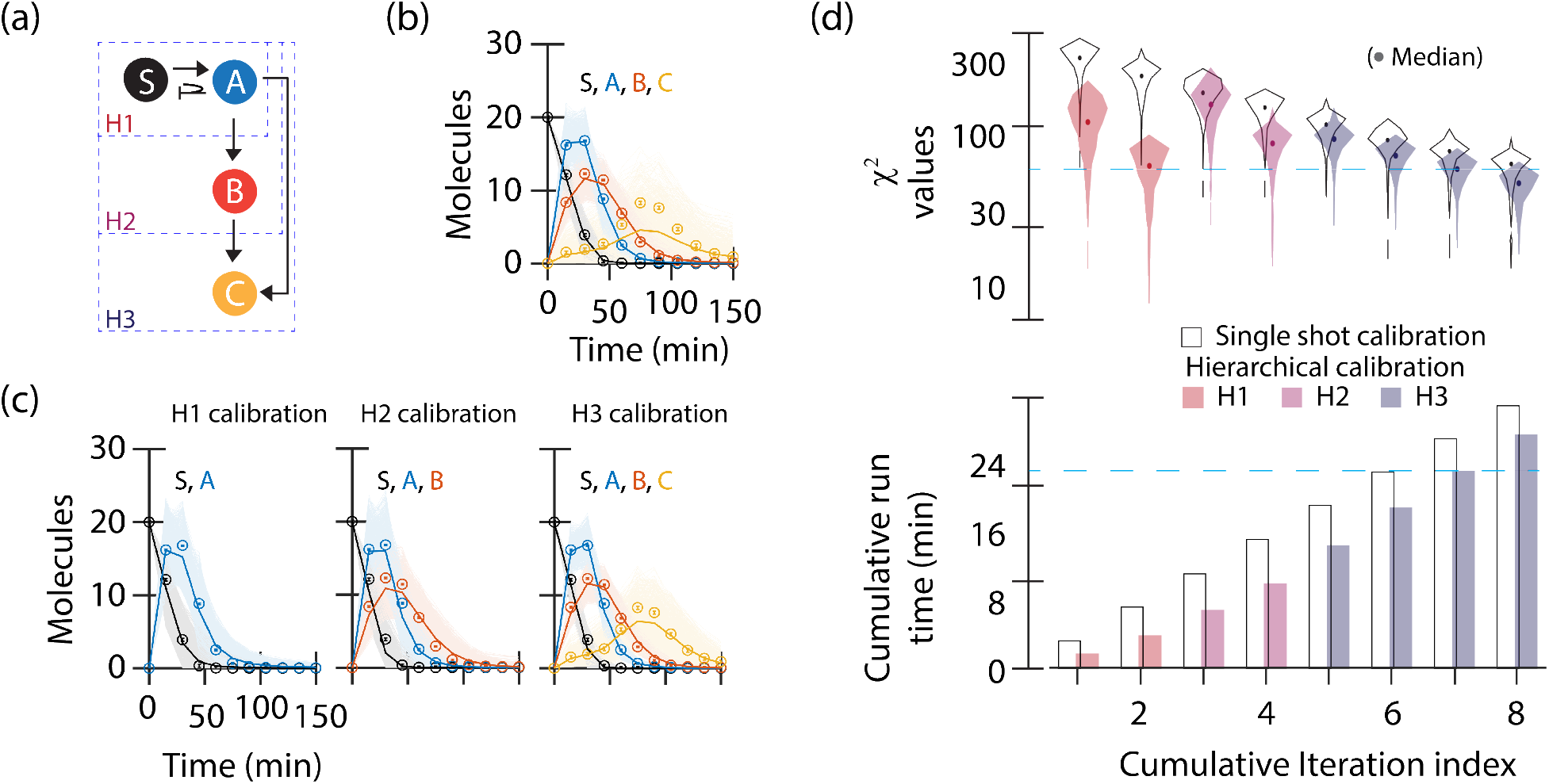
Exploiting model structure for more efficient model fitting. (a) Schematic of the Feed-forward loop (FFL) system. (b) Model fitting due to oCal strategy is shown. (c) Model fitting after H1 (left), H2 (middle) and H3 (right) are shown. In subfigs b and c, thin lines represent model predictions due to individual parameter combinations selected at the final iteration, the thick lines track the mean of predictions over selected parameter combinations and open circles represent the data used for fitting. (d) Efficiency of oCal and hCal strategies are compared in terms of accuracy (top) and time complexity (bottom).

**Figure S8:**
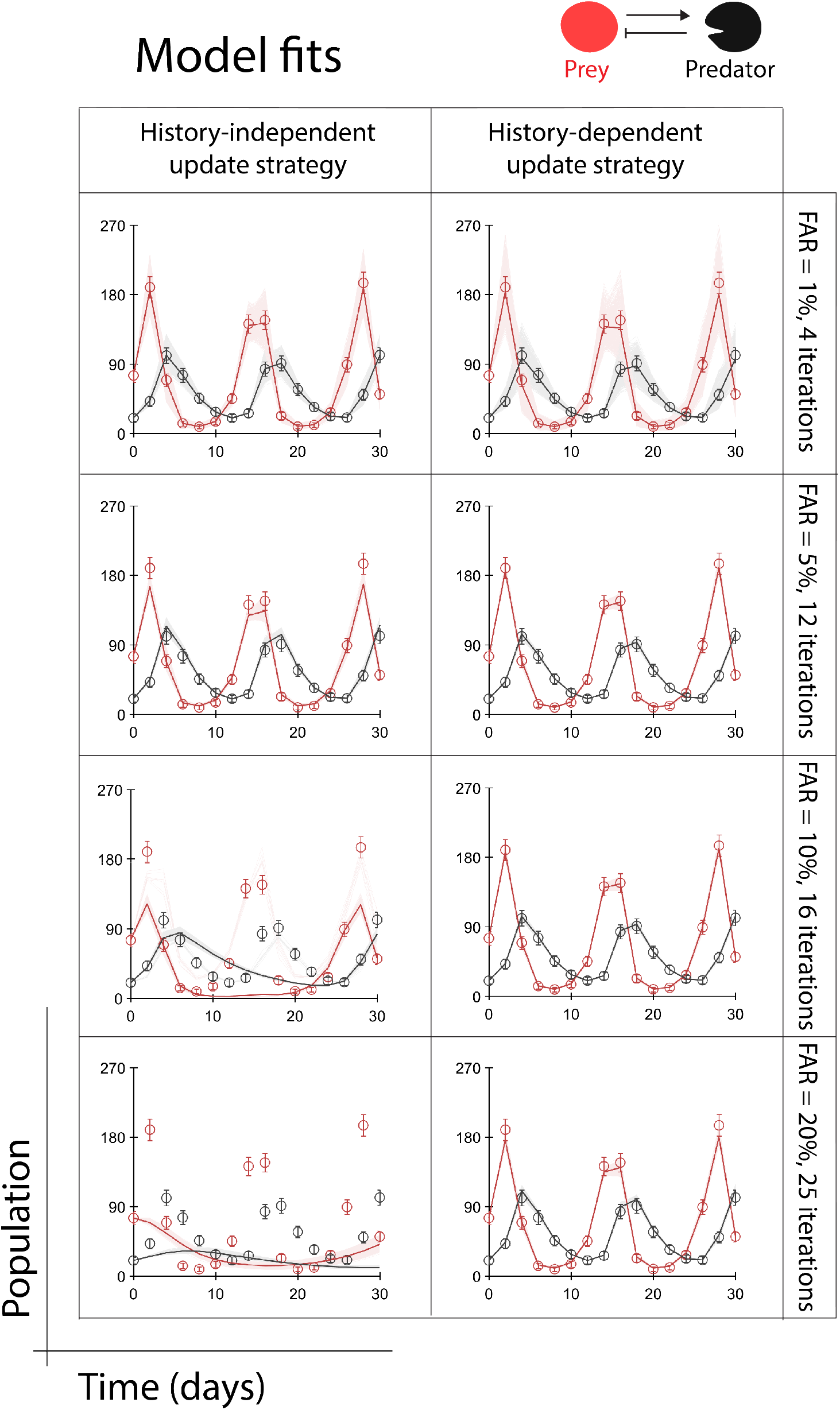
Tuning algorithmic options. Here we see how the Lotka-Volterra model fitting is affected when we vary the update strategies (History-independent update strategy and History-dependent update strategy) and FAR values. Here the thin lines represent model predictions due to individual parameter combinations selected at the final iteration, the thick lines track the mean of predictions over selected parameter combinations and open circles represent the data used for fitting.

**Figure S9:**
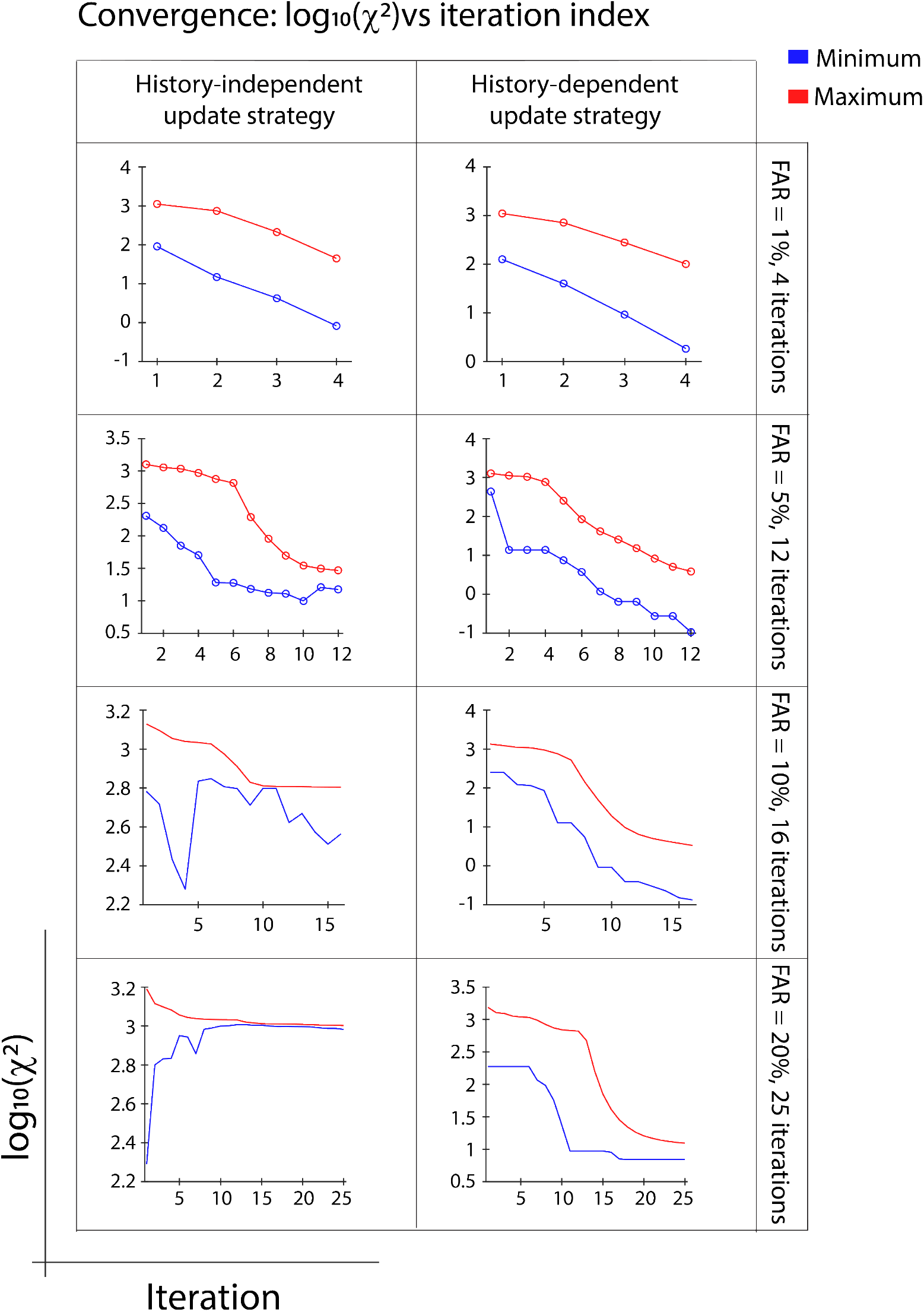
Tuning algorithmic options. Here we see how the Lotka-Volterra model fitting is affected when we vary the update strategies (History-independent update strategy and History-dependent update strategy) and FAR values. The minimum (blue) and the maximum (red) of the χ^2^ values corresponding to the parameter combinations selected in every iteration is plotted.

**Figure S10:**
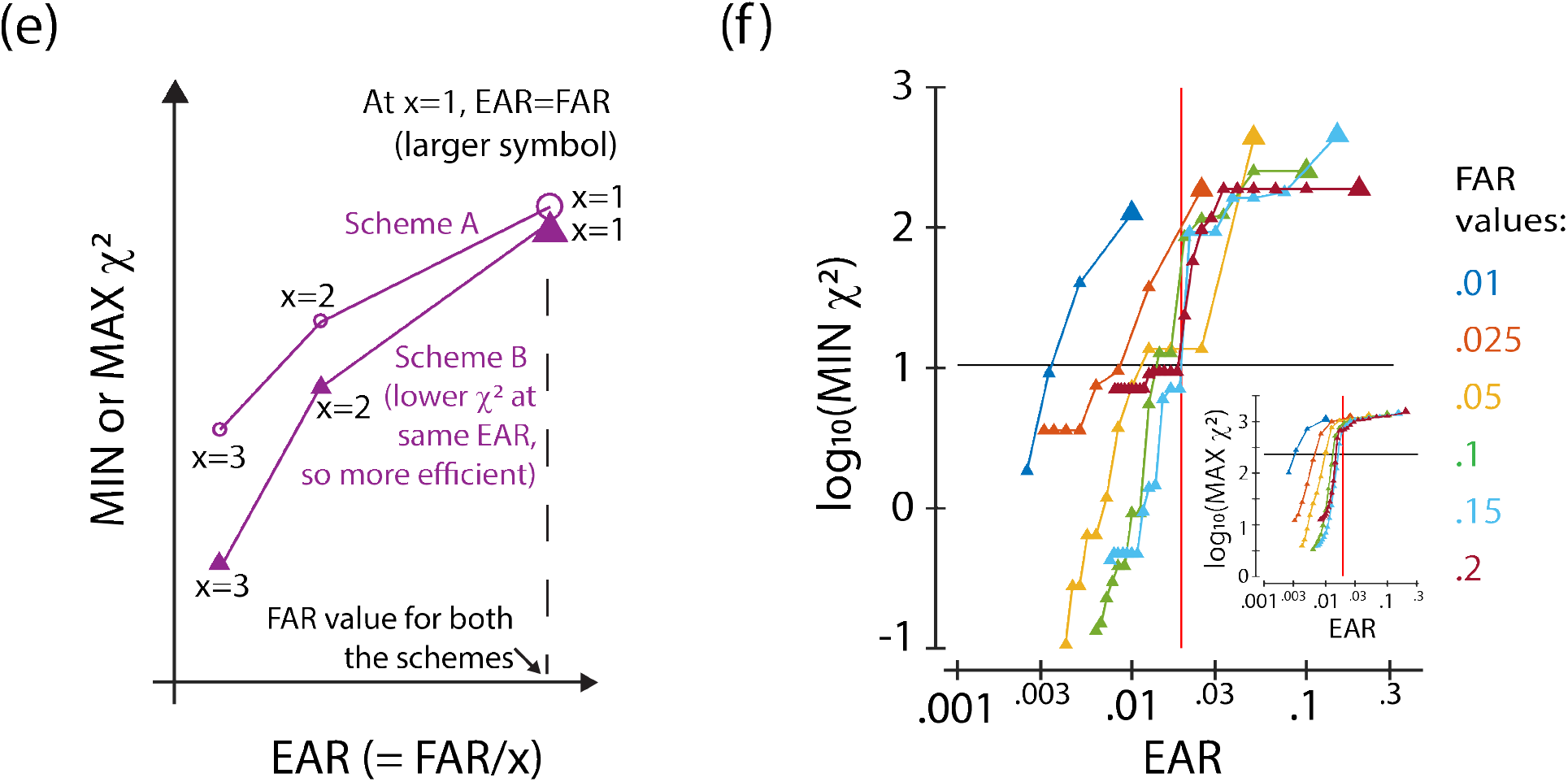
Accuracy and efficiency of the parameter estimation algorithm. (e) To characterize the efficiency of the estimation process we look at accuracy and computational cost. Accuracy is inferred by the minimum of χ^2^ values corresponding to the selected combinations; mean or maximum value can also be considered. Computational cost is measured by the total number of parameter combinations sampled and simulated. Thus the cost is inversely proportional to Effective Acceptance Rate (EAR), as the number of parameter combinations selected in each iteration is kept constant throughout the analysis. An efficient process would have a lower χ^2^ statistic at a given EAR value, or high EAR when we fix the χ^2^ statistic. (f) We characterize the efficiency for each iteration of a parameter estimation process to fit the Lotka Volterra model to computationally generated data, using ABC-FAR with different values of FAR. We see that χ^2^ statistic initially decreases with decrease in EAR (faster for higher FAR value) and then saturates (at higher EAR for higher FAR value). The efficiency of our algorithm compares well with that reported for ABC-SMC, shown in the figure by the intersection of black and red solid lines.

**Figure S11:**
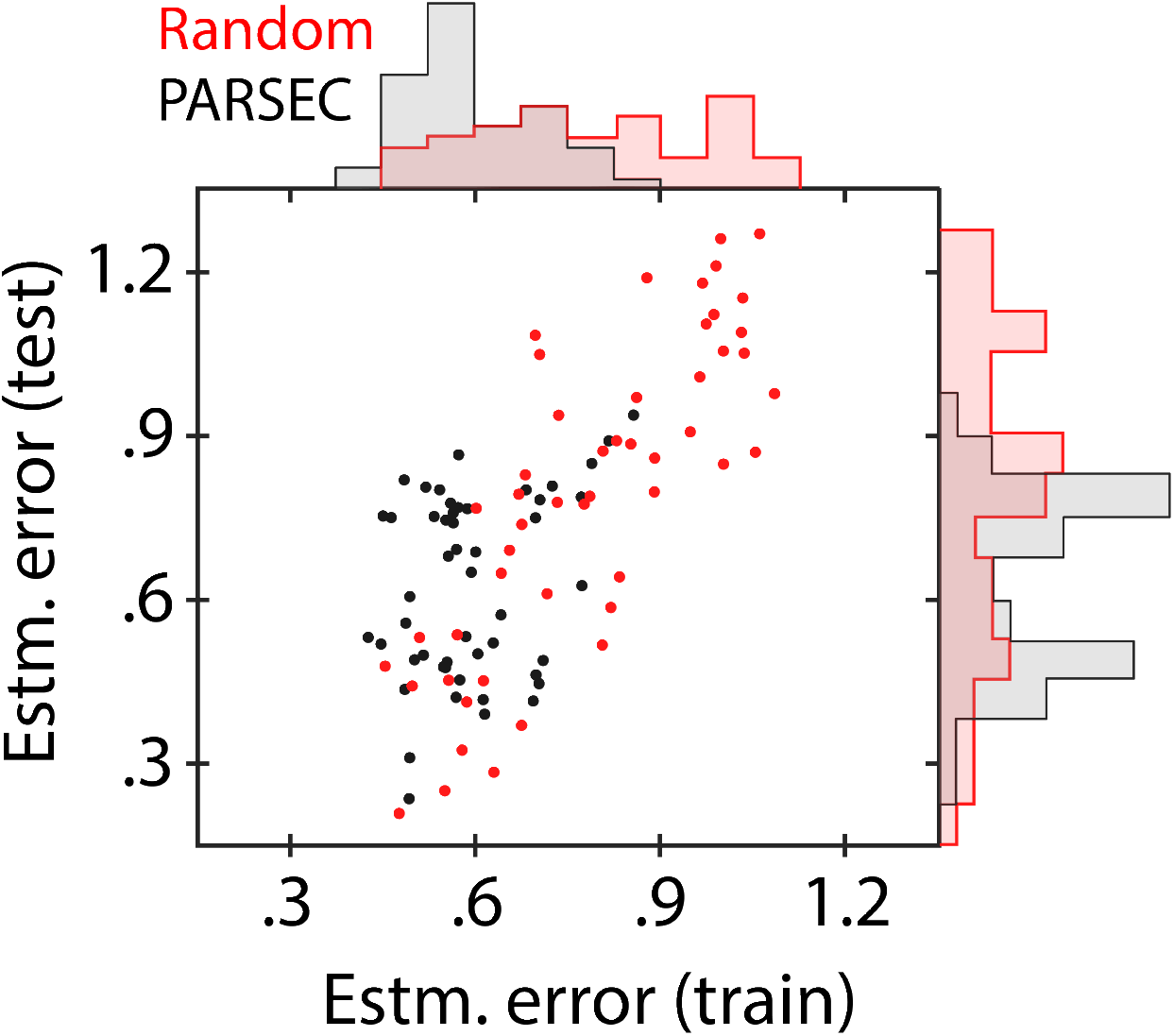
Robustness of PARSEC(k) (b) Here we consider a nine-fold uncertainty characterized by 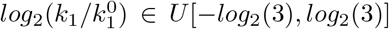 and 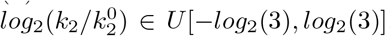. where U[a, b] denotes a uniform distribution bounded by a and b. PSI are evaluated at nine training samples (Θ^*k*^) representing this distribution, are used to construct the PARSEC-PSI vectors. These are grouped using k-means clustering to inform the selection of PARSEC(k) designs. 100 PARSEC(k) and 100 random designs are evaluated for estimation accuracy at the training samples (Training analysis) and at four new test samples (Ω^*k*^, test analysis). Sampling was done via Latin Hypercube Sampling. The average estimation error across the training samples and the test samples are evaluated as train and test estimation errors respectively. The corresponding marginals are also shown. The plot highlights that PARSEC(k) designs are more informative. The 2D-plot of the train and test estimation errors verify the equivalence of performance of the best few PARSEC(k) designs against stochastic sampling, indicating their robust performance.

**Figure S12:**
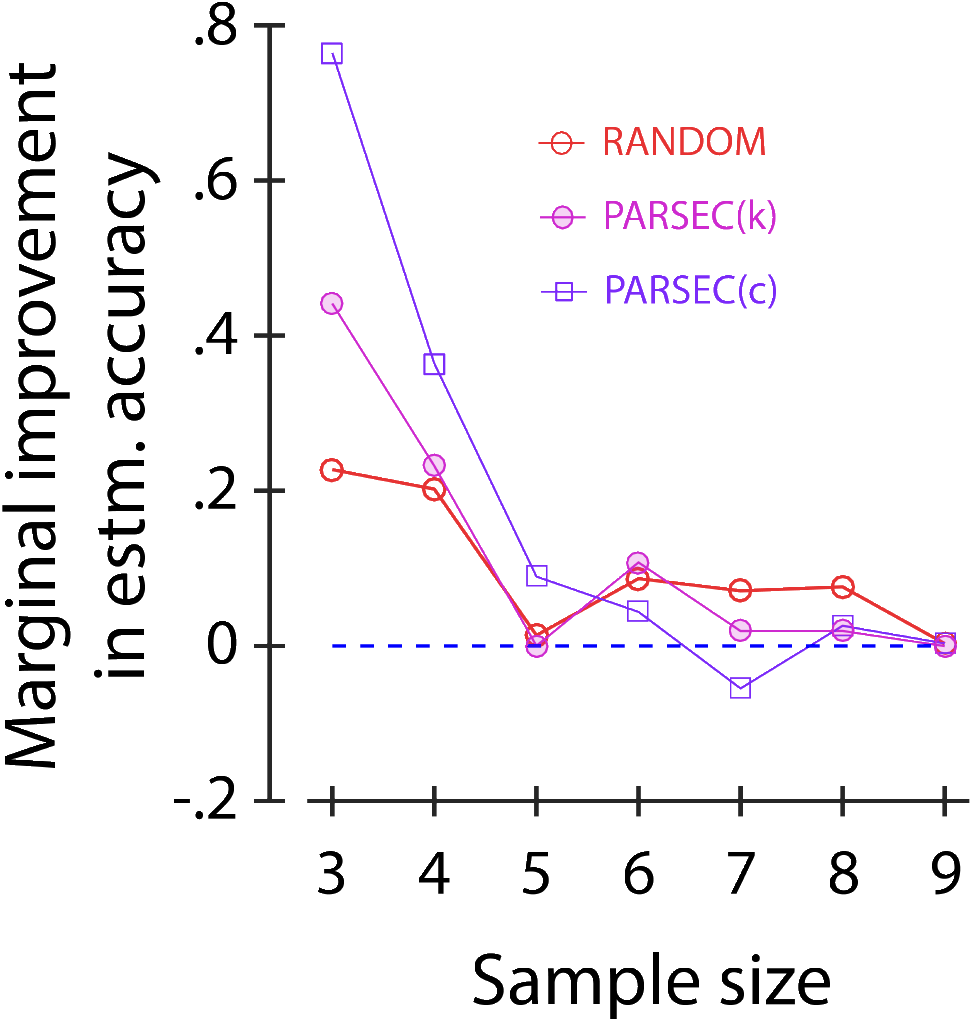
Marginal gain in estimation accuracy due to increase in experiment sample size. (a) We plot the dynamics of the three-gene repressilator system for the parameter combination used for the design. (b) One of the realizations of c-means clustering of the measurement candidates for cluster multiplicity ranging from two to nine is shown. Clusters are identified using different colors.

**Figure S13:**
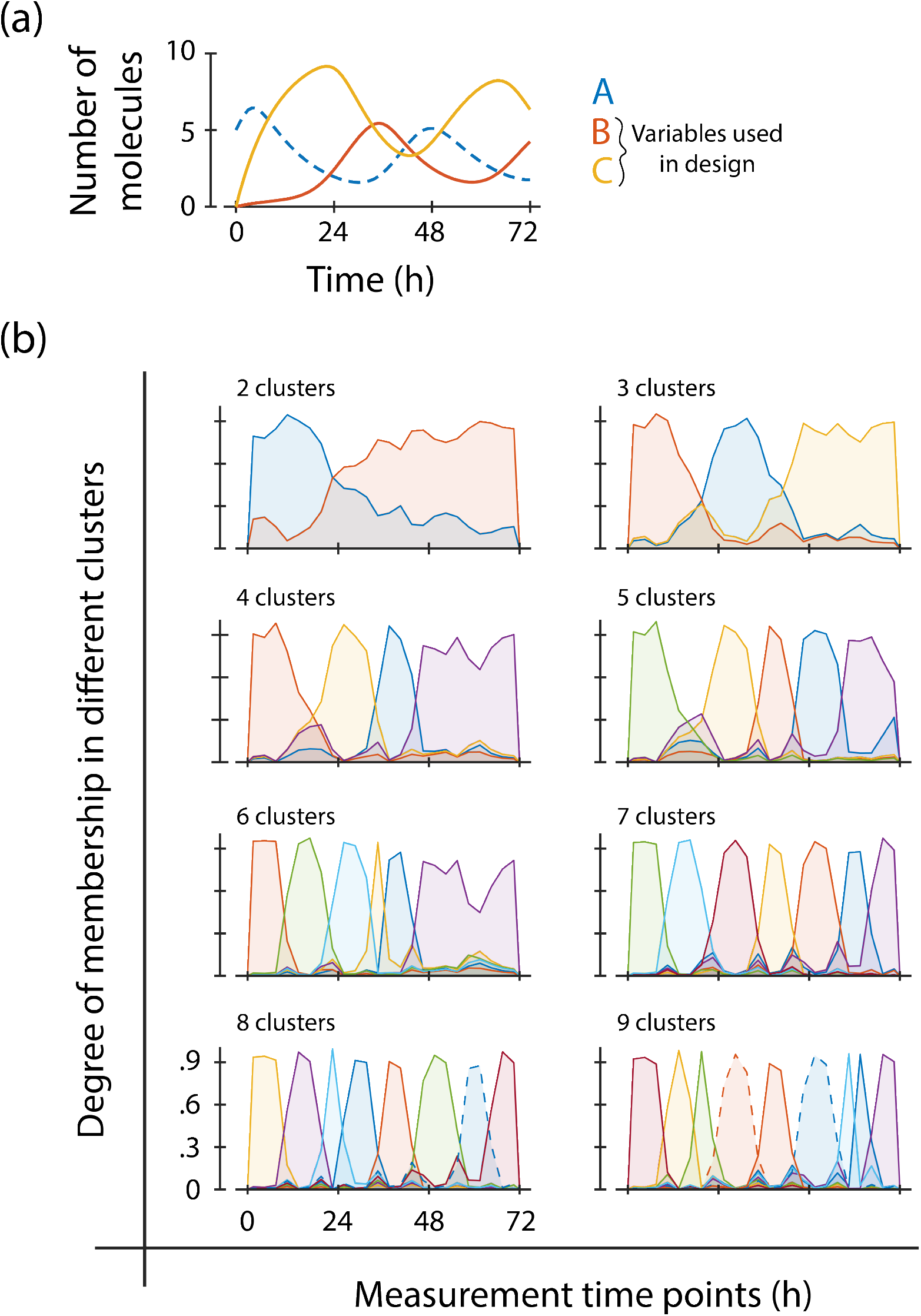
Partitioning via c-means clustering algorithm. (a) We plot the dynamics of the three-gene repressilator system for the parameter combination used for the design. (b) One of the realizations of c-means clustering of the measurement candidates for cluster multiplicity ranging from two to nine is shown. Clusters are identified using different colors.

**Figure S14:**
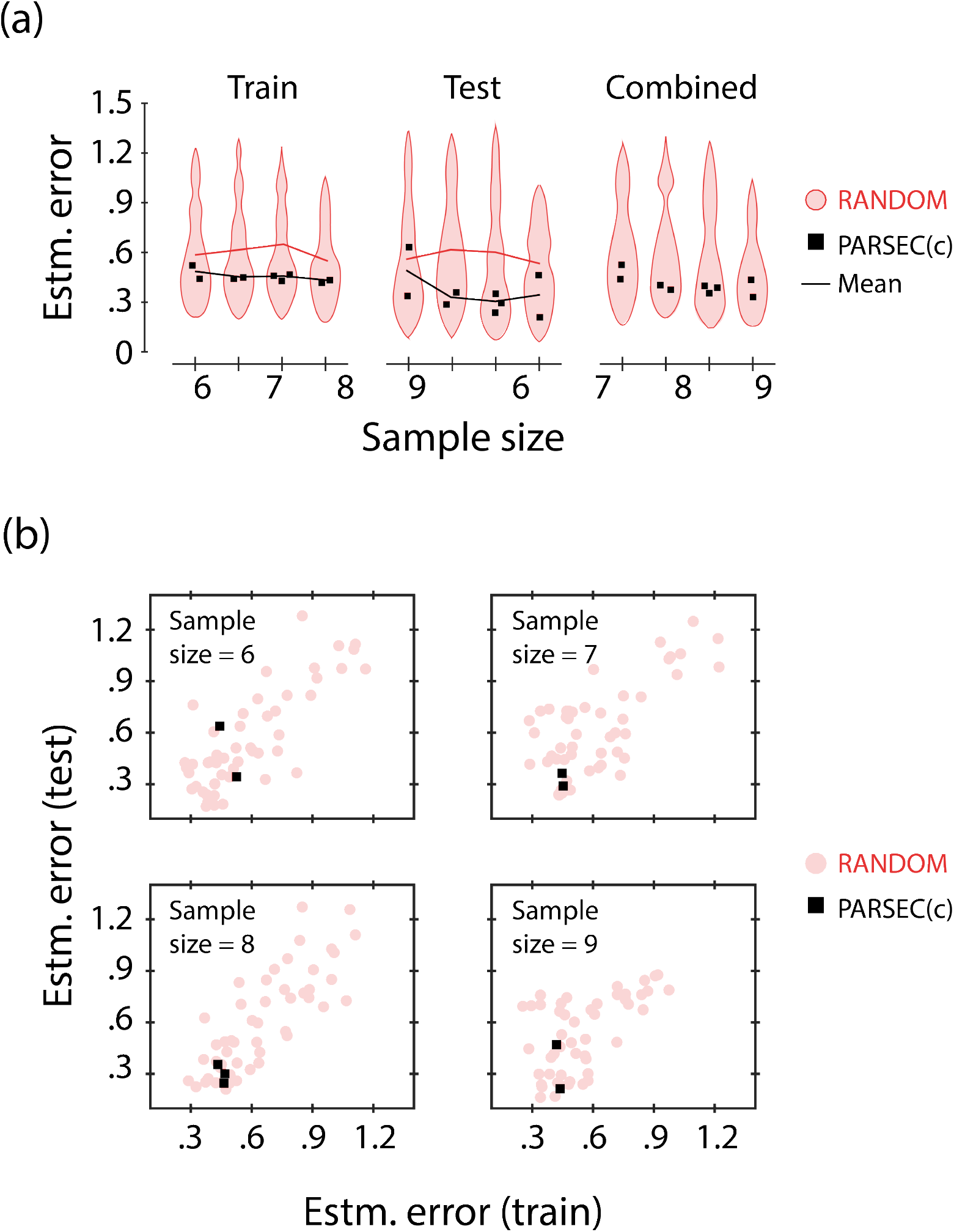
Performance of generalist design constructed via PARSEC(c) We use PARSEC(c), which uses c-means clustering, to efficiently identify informative generalist designs according to the specifications used for the analysis summarized in Figure 5 (Main text). We also consider the same training and test samples as used there. The corresponding dynamics and a realization of the c-means clustering is shown in Supplementary Figure S15. (a) We plot the distribution of estimation error averaged across the training (Train statistics, left) and testing (Test statistics, right) samples for 50 random designs (red), and the estimation error for each of the unique PARSEC(c) designs (black) considered in the analysis. We also show the average statistics for the estimation error for the two modalities of design as a function of sample size. The average estimation error reduces with an increase in sample size. Although PARSEC(c) may miss the most informative designs, on average it identifies designs more informative compared to random designs with about 44 times fewer calculations. (b) The 2D-plot of the train and test estimation errors verify the equivalence of performance of PARSEC(c) against stochastic sampling, indicating their robust performance.

**Figure S15:**
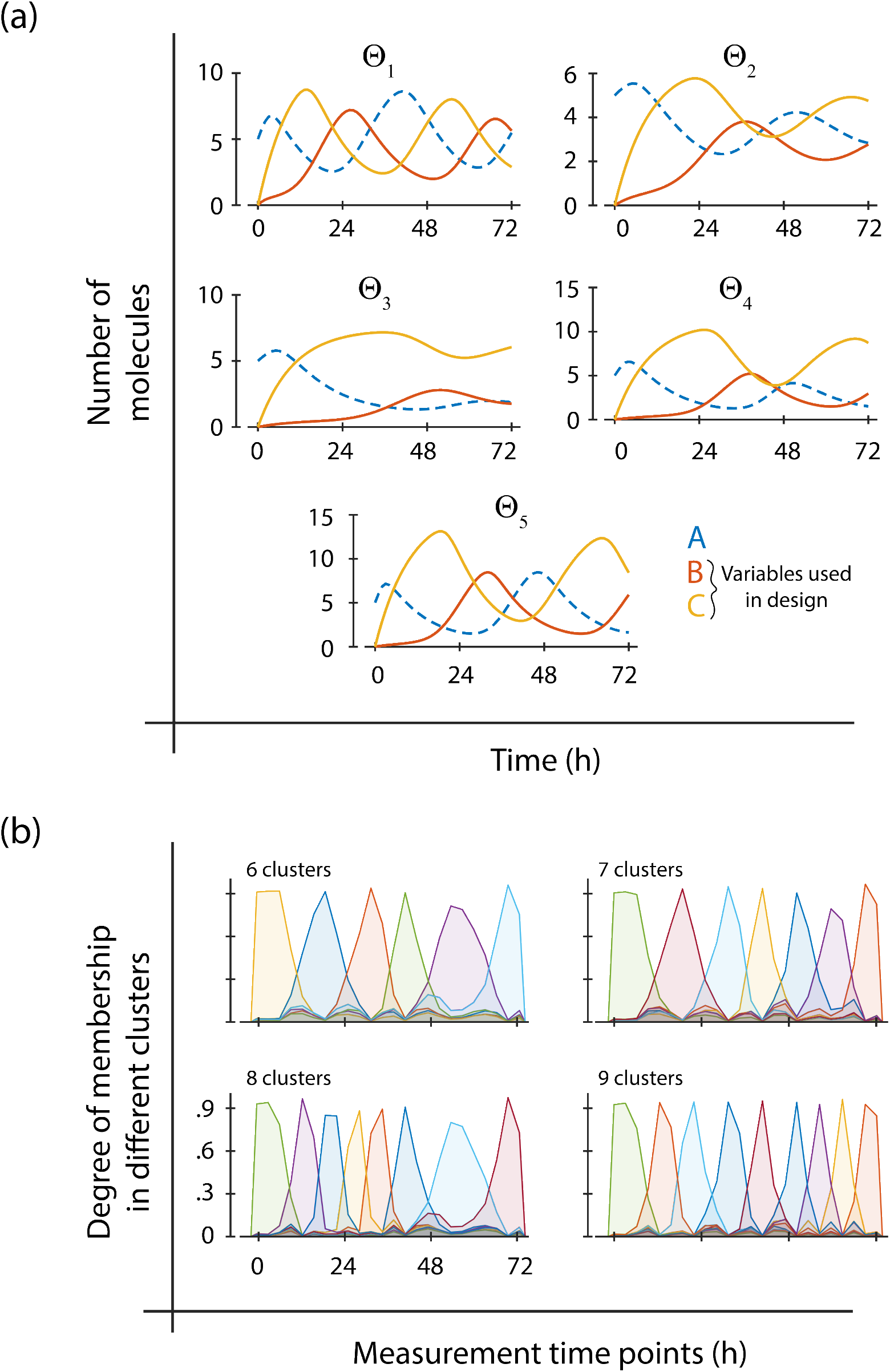
Dynamics and c-means clustering involved in constructing generalist designs using PARSEC(c) (a) We plot the dynamics of the three-gene repressilator system for the five training samples used in the design construction. (b) One of the realizations of c-means clustering of the measurement candidates for cluster multiplicity ranging from six to nine is shown. Clusters are identified using different colors.

## Notes

### Competing Interest Statement

The authors have declared no competing interest.

